# Computational Repurposing of FDA Approved Drugs Against Cyclin Dependent Kinase (CDK2)

**DOI:** 10.64898/2026.10.27.762194

**Authors:** Girija Deevi, Bhupender Singh

**Affiliations:** Biological E limited, Hyderabad, Telangana, India; CRO Scientist Biotechnika Info Labs Pvt Ltd

**Keywords:** Drug Repurposing, CDK2, Swiss ADME, FDA-approved drugs, PLIP

## Abstract

*Drug repurposing* is an approach for identifying new therapeutic with existing FDA-approved drugs. Cyclin-dependent kinase 2 (CDK2) is an important regulator of cell-cycle progression. The present study aimed to identify FDA-approved drugs with potential binding activity against CDK2 using a Computational drug-repurposing approach. The crystal structure of Human CDK2 (PDB ID: 3PY0) was used for Molecular docking, followed by Protein–Ligand Interaction Profiler (PLIP) analysis to evaluate Protein–Ligand interactions. Further for drug-likeness and pharmacokinetic properties using Swiss ADME and for Predicted toxicity using ProTox Was done. Based on this docking affinity, interaction profiles, drug-likeness, pharmacokinetics and toxicity, the best target was selected. Carteolol demonstrated the most balanced overall computational profile and was selected as the primary candidate for potential computational finding to provide repurposing against CDK2. Experimental studies are required to validate its direct binding and inhibitory activity against CDK2.

## Introduction

New drug development involves identification of suitable biological targets, Analysis and preclinical studies and clinical evaluation. Despite the use of several resources many drugs fail during development and do not provide extensive results due to toxicity or pharmacokinetic properties. So, because of these reasons researchers started exploring to make new drug developments and its applications more efficient*. **Drug repurposing*** is also referred to as drug repositioning, in which an existing drug is investigated for a therapeutic application different from its original indication (Pushpakom *et al.,* 2019). It’s not like conventional drug discovery where chemical moiety is developed for a particular target. This drug repurposing actually starts preliminary with a compound where its information is already available like chemical properties, pharmacologic, kinetics, toxicity and clinical data. These early stages of drug development may shorten the time which is required to identify potential therapeutic candidates (Pushpakom *et al.,* 2019). Importantly this Drug repurposing can be done and reach to a certain approach by systematic experimental and computational investigations. With help of large set of data bases,chemical libraries and Insilco-Methods,computational biology .Computational methods helps to identify binding and relationships with existing drugs in the data bases with molecular targets .In this way it helps researchers to examine large numbers of compounds and prioritize candidates for subsequent experimental investigation (Polamreddy & Gattu *et al.,*2019).A drug which have its effectiveness for its original may have its own mechanism of action that could be different or not relate to this new target .Therefore its interaction with new target with all parameters, structure, pharmacokinetic behaviour and toxicity must still be investigated. (Pushpakom *et al.,*2019).

## Cyclin-Dependent Kinase 2 (CDK2)

Cyclin-dependent kinase 2 (CDK2) is a serine/threonine protein kinase which is cyclin-dependent kinase family. They are important cell proliferation regulators as they help in coordination of cell progression through different stages of the cell cycle. Activity and regulation by cyclin proteins which helps in formation of active cyclin–CDK complexes. Regulation of the transition from the G1 phase to the S phase is guarded by CDK2 and involved in DNA replication (Knudsen *et al.,* 2025; Scheidt *et al.,* 2025). The Cell cycle has four major stages G1, S, G2 and M. During G1, the cell grows and prepares for DNA synthesis. During S phase duplication of the cellular DNA happens in G2 cell for mitosis and the M phase involves chromosome segregation and cell division. Process happens correctly and sequentially to ensure DNA replication and cell division. CDK2 mainly plays a role in the G1/S transition thus promoting the cell into the DNA-synthesis phase (Knudsen *et al.,* 2025). Cyclin E and cyclin A should be closely associated otherwise CDK2 does not function independently. Cyclin E levels increase and cyclin E with CDK2 produces an active kinase complex during late G1. Phosphorylation of proteins happens due to this complex which is involved in cell-cycle progression and creates a condition to enter S phase. G1/S transition completed cyclin A– CDK2 complexes become important during the S phase where DNA replication and cell-cycle progression occurs. Conformation changes occur with cyclin binding which activates kinase phosphorylation and endogenous CDK inhibitors provide additional levels. This regulation provides correct and on time activity of CDK2 during stages of cell cycle. Any changes or deviation from this leads to abnormal cellular proliferation (Knudsen *et al.,*2025)

## CDK2 and Cancer

There is a relationship between CDK2 and cancer which is uncontrolled cell growth, a main characteristic of cancer. Any changes in cyclin–CDK Signalling can disturb normal cell-cycle checkpoints and cycle which allows cells to rapidly divide. Same to this cyclin E activity and alterations in the cyclin E–CDK2 pathway leads to different cancers have contributed to interest in CDK2 as a potential therapeutic target (Scheidt *et al.,* 2025).

## Structural Relevance of CDK2

In this study human CDK2 was selected as the molecular target; the crystal structure represented by PDB ID 3PY0 was used as the structural basis for the docking investigation. 3PY0 has CDK2 in complex with the ligand SU9516 which provides 3D experimental representation of the target which is useful to study structure-based analysis (Betzi *et al.,* 2011). Due to its biological importance in cell-cycle regulation and its potential therapeutic target CDK2 selection was done. Ligand-bound experimental structure such as 3PY0 further enabled the investigation of whether existing FDA-approved drugs could demonstrate predicted interactions with the CDK2 binding region.

## FDA-Approved Drugs

FDA-approved drugs are reviewed and authorized by the U.S. Food and Drug Administration (FDA) for particular use after careful evaluation of their quality, Clinical studies, drug efficacy, and manufacturing standards. These drugs may include small-molecule compounds and biological products developed to prevent, diagnose or treat particular diseases. For drug repurposing FDA-Approved Drugs are a good source because they are being monitored for safety, pharmaceutical development for their original indications, affects, safety issues. Using these drugs with known and well-studied descriptions one can reduce time, cost, risk associated with discovering completely new chemical Structures (Pushpakom *et al.,* 2019). FDA approval does not mean a drug will be effective against a different disease or molecular target. Repurposing study Should be performed with additional studies to determine ligand and receptor interaction, whether drug is reaching its expectation with target. It is possible with Computational screening of FDA-approved compounds. This Computational screening will check binding before experimental validation.

## Computational Drug Repurposing

Computational drug repurposing uses bioinformatics tools and structure-based ways to find new potential for existing drugs. Techniques Include molecular docking, virtual screening, molecular similarity analysis and pharmacophore-based methods can be used for drug–target relationships before experiment Validation. This Method is particularly used for large compound library screening and targeting candidates for further investigation

## Method

### Materials and Computational Resources

- UniProt
- RCSB Protein Data Bank (PDB)
- ZINC20
- PyMOL
- PDB2PQR
- Webina/AutoDock Vina
- PLIP
- SwissADME
- ProTox-3

### Target Characterization and Library Preparation

Cyclin-dependent kinase 2 (CDK2) was selected as the molecular target for the present drug-Repurposing Study.UniProt was used as the primary protein sequence and functional annotation database. The human CDK2 protein entry was identified using the protein name and organism information. *UniProt* provides information including the gene name, protein name, amino-acid sequence, protein length, biological function and other relevant functional annotations. The UniProtKB accession number is P24941, corresponding to a reviewed protein entry with a length of 298 amino acids. The target information and protein sequence were retrieved from the UniProt database (UniProt Consortium, 2025).

Reference: https://www.uniprot.org/uniprot/P24941

**Table 1:** Details of Human cyclin-dependent kinase 2 (CDK2) using the UniProtKB/Swiss-Prot database (P24941).

| **Parameter** | **CDK2 information** |
|---|---|
| Protein | Cyclin-dependent kinase 2 |
| Gene | CDK2 |
| UniProt accession | P24941 |
| UniProt entry | CDK2_HUMAN |
| Organism | *Homo sapiens* |
| Length | 298 amino acids |
| Molecular mass | 33,930 Da |
| Annotation score | 5/5 |

**Figure 1:**
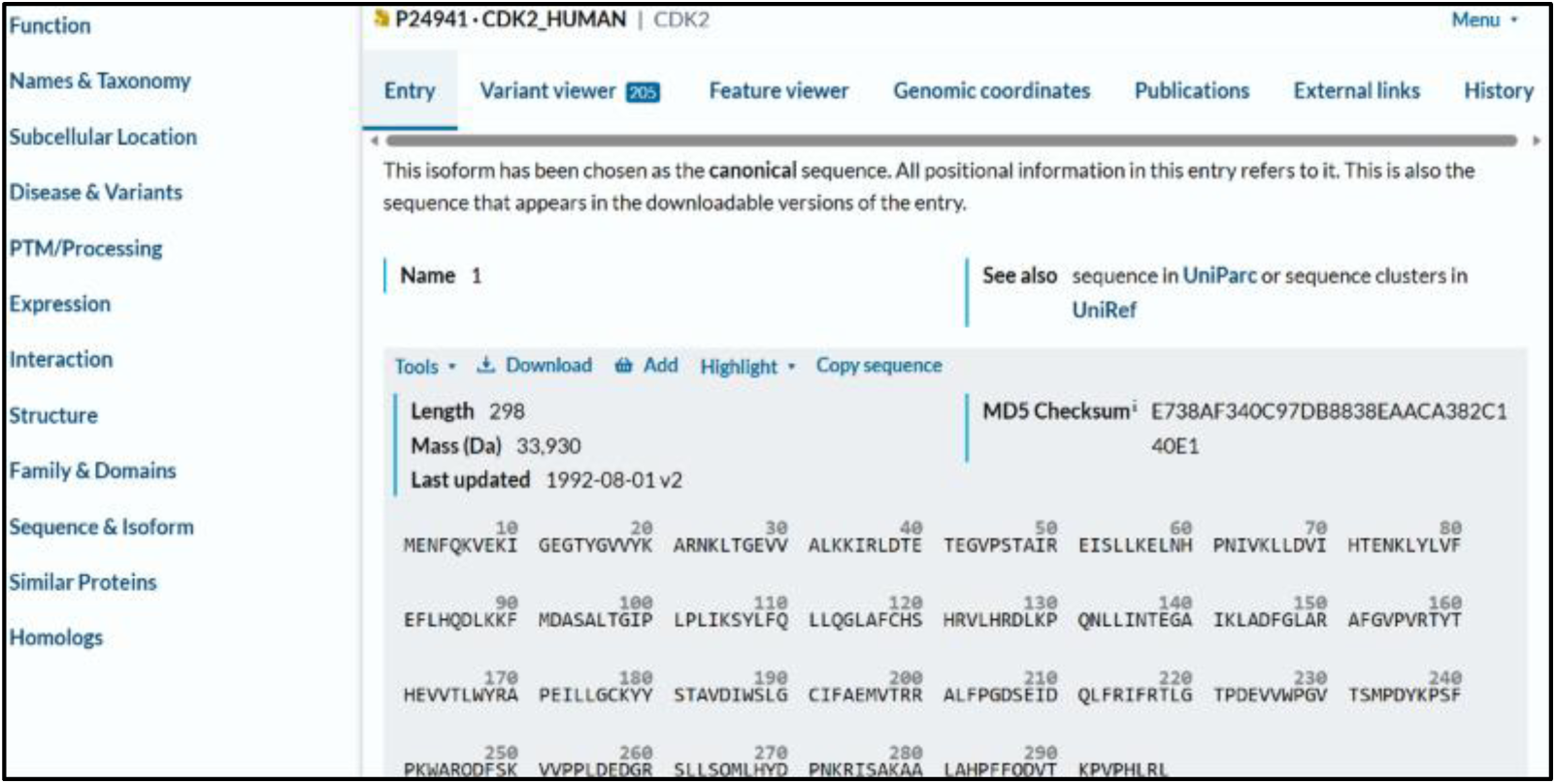
Accession P24941 (CDK2_HUMAN) sequence was obtained from the UniProtKB

Before moving on to three-dimensional structural analysis, the identity and fundamental sequence properties of human CDK2 were verified using the UniProtKB/Swiss-Prot database. A suitable receptor structure for molecular docking was then chosen by looking at the corresponding experimental structures that were available in the RCSB Protein Data Bank

### Retrieval and Selection of CDK2 Crystal Structure

3D structure of the protein was subsequently required. The RCSB Protein Data Bank (PDB) (https://www.rcsb.org/)provides structural records of Human CDK2 that gives experimentally determined three-dimensional structures of proteins and protein–ligand complexes help to select a receptor for docking studies.

The CDK2 was searched as target protein corresponding to Homo sapiens. Among all, PDB ID 3PY0 was selected for the present study. The structure was determined using X-ray diffraction and human CDK2 in complex with the co-crystallized small-molecule ligand SU9516.This provides an experiment of human CDK2 with co-crystallized ligand within the relevant binding region. Bound ligand will provide structural reference for the docking region and prepare the receptor for screening of FDA-approved compounds. The PDB structure was downloaded in PDB format. The structural information obtained from 3PY0 was subsequently used to prepare the CDK2 receptor for molecular docking of the FDA-approved drug library.

**Figure 2:**
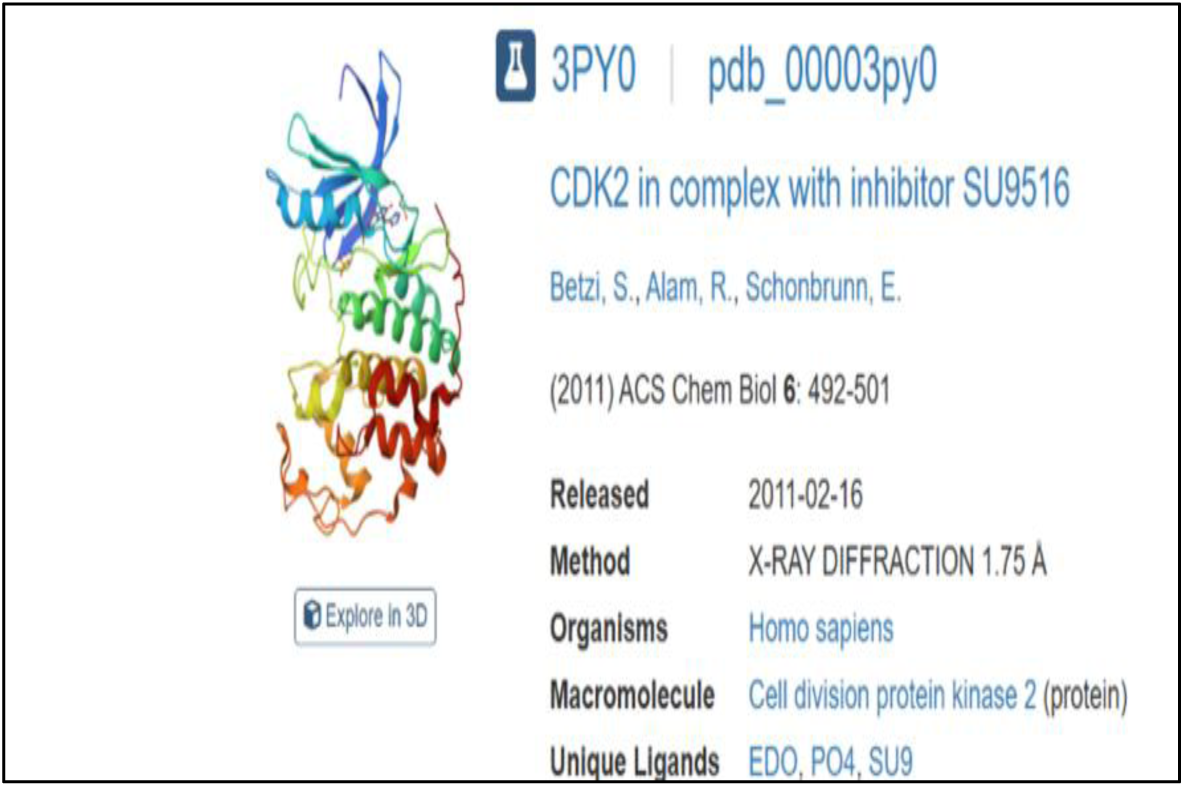
Human CDK2 structure From RCSB Protein Data Bank

**Figure 3.**
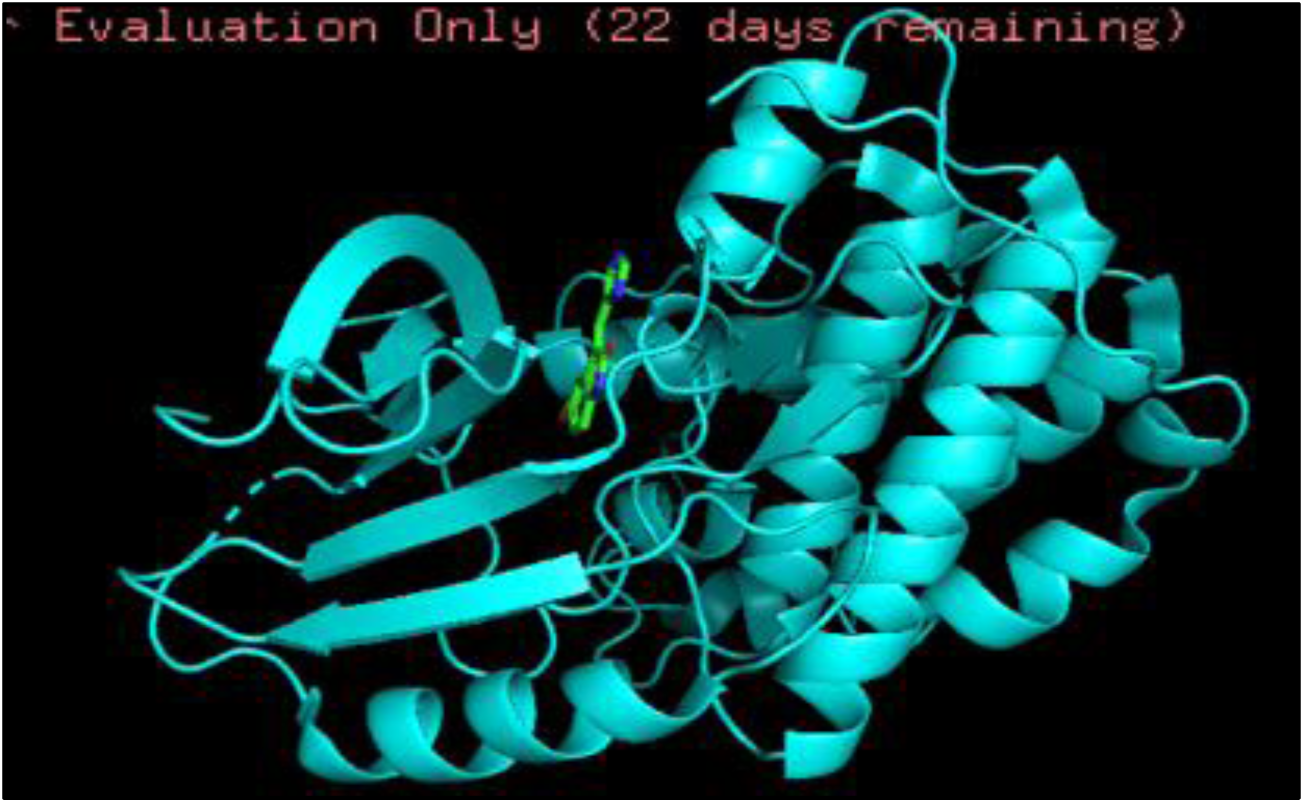
: Human CDK2 (PDB ID 3PY0) showing the co-crystallized inhibitor SU9516 within the ligand-binding region 3D structure

### Protein Structure Preparation

The determined CDK2 structure, PDB ID 3PY0 downloaded from the RCSB Protein Data Bank in PDB format and used for receptor preparation. The downloaded structure was first processed using the PDB2PQR (https://server.poissonboltzmann.org/pdb2pqr) web server to generate a computationally prepared representation of the protein. PDB2PQR was configured to use PROPKA for protonation-state assignment at pH 7.4. The prepared structure in PQR format.

PDB2PQR calculation, the generated PQR file and output files were downloaded. The co-crystallized inhibitor SU9516 (SU9) was retained during the initial protein-preparation stage because it was used as a structural reference for the binding region. Before molecular docking of the FDA-approved drugs, SU9516 was removed from the receptor to prevent competition with the docked ligands. Solvents, Hetero atoms and other unwanted crystallographic components were also removed during the final step using Pymol. The resulting cleaned CDK2 structure was saved and used as the receptor for molecular docking.

**Figure 4:**
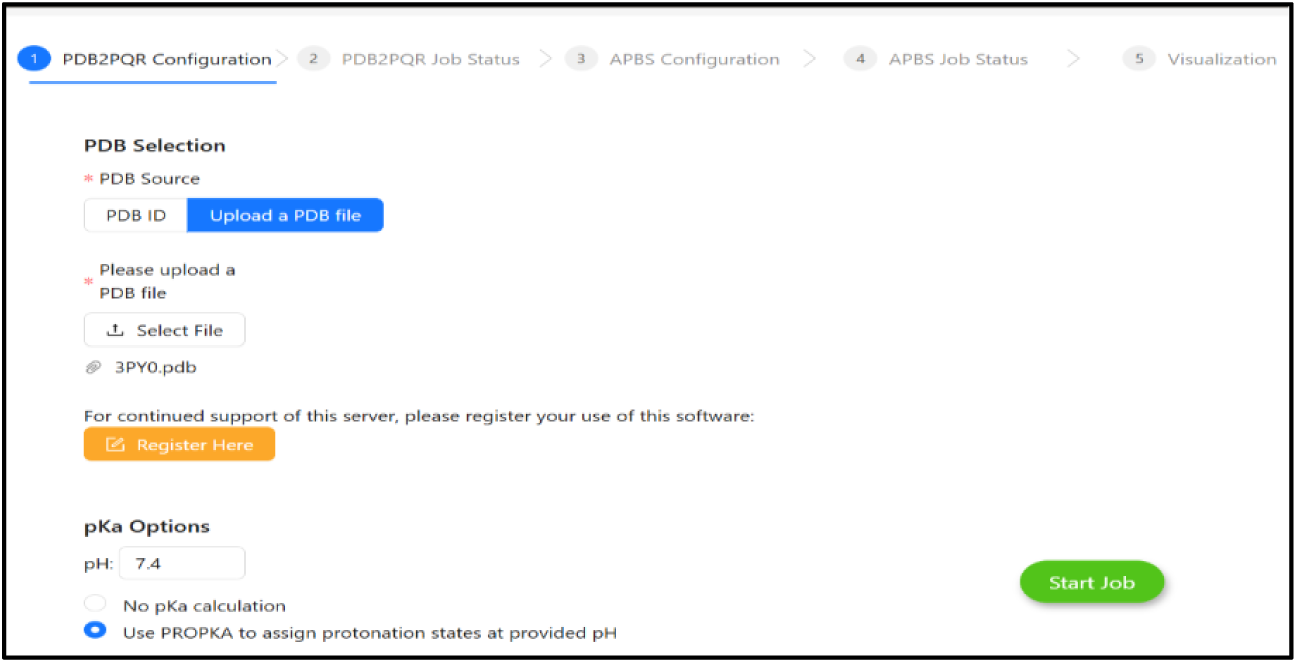
Human CDK2 structure (PDB ID 3PY0). Preparation using PDB2PQ

**Figure 5:**
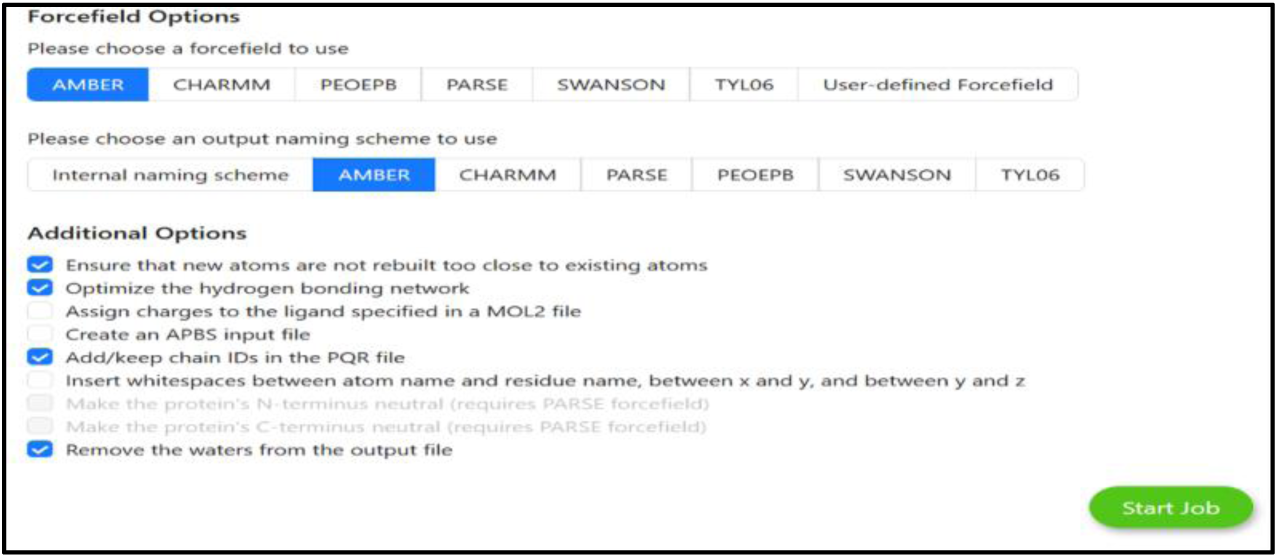
Human CDK2 structure (PDB ID 3PY0) preparation using PDB2PQR with selected options

### FDA-Approved Drug Library Preparation Using ZINC20

After preparation of the CDK2 receptor. The ZINC20 database(https://zinc20.docking.org/) was selected as the source of the ligand library because it provides access to commercially available and biologically relevant small molecules, including collections of approved drugs suitable for virtual screening. The FDA-approved drug selected from ZINC20.The compounds were downloaded in a structured format compatible with the docking workflow.

**Figure 6:**
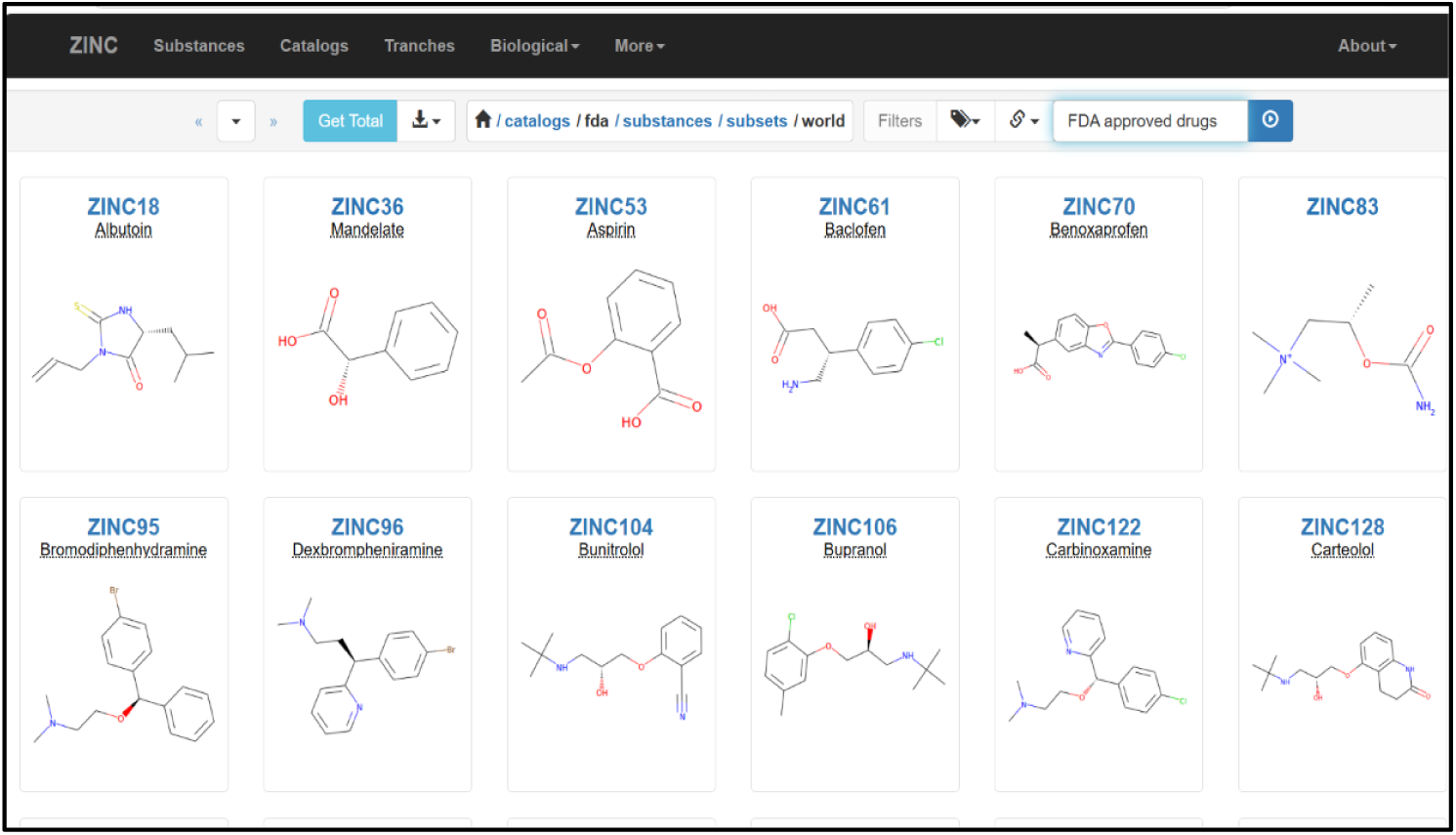
Showing FDA-approved drug from ZINC20

The collected drug information was organized into a ligand library prior to molecular docking. The drug name, ZINC identifier, and corresponding SMILES notation were documented for each compound throughout the computational screening workflow.

## High-Throughput Virtual Screening and Molecular Docking

### Receptor Preparation and Docking Using AutoDock-Vina PrepServer

Following protein preparation, the human CDK2 structure (PDB ID: 3PY0) was refined using the AutoDock-Vina PrepServer to generate a receptor for molecular docking. The **3PY0** structure contained several crystallographic non-protein components that were identified during the receptor-preparation step. The detected HETATM components included EDO, HOH, PO4, and the Co-crystallized inhibitor SU9516 (SU9). These components were selected for removal to prevent non-essential crystallographic molecules and the original inhibitor from interfering with the docking of the selected FDA-approved compounds.

**Figure 7:**
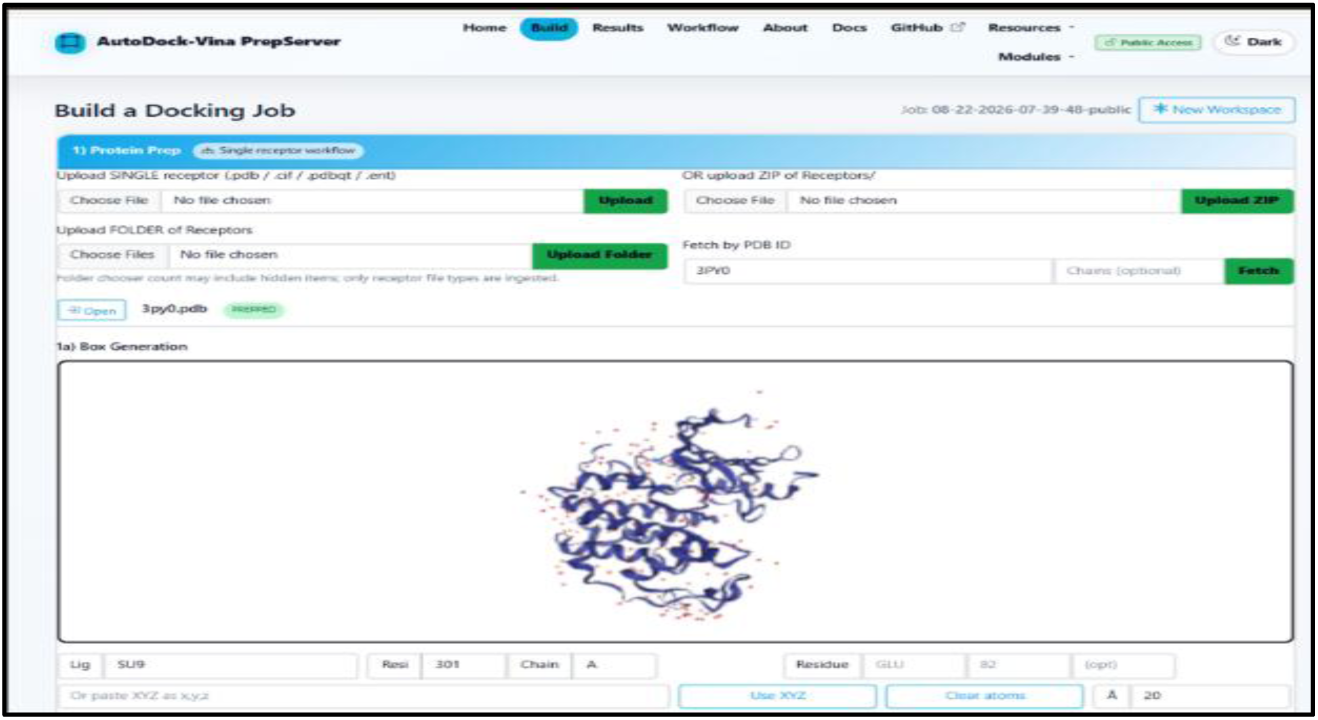
Auto Dock-Vina workflow showing receptor preparation and binding site

The binding region was defined using the position of the co-crystallized SU9516 ligand before its removal. The receptor was processed by the *Auto Dock-Vina PrepServer* and converted into PDBQT format, which was subsequently used for molecular docking the binding region was defined using the position of the co-crystallized SU9516 ligand before its removal. The receptor was processed by the *Auto Dock-Vina PrepServer (*https://www.autodockvina.com/build)and converted into PDBQT format, which was subsequently used for molecular docking

**Table 2:** Software’s used for docking

| **Software/Tool** | **Version** | **Purpose** |
|---|---|---|
| AutoDock-Vina PrepServer | Web-based | Receptor and ligand preparation |
| Webina | 1.0.5 | Molecular docking |
| AutoDock Vina | 1.1.2 | Docking engine underlying Webina |
| PyMOL(TM) 3.1.8 | 3.1.8 | visualization |

**Figure 8:**
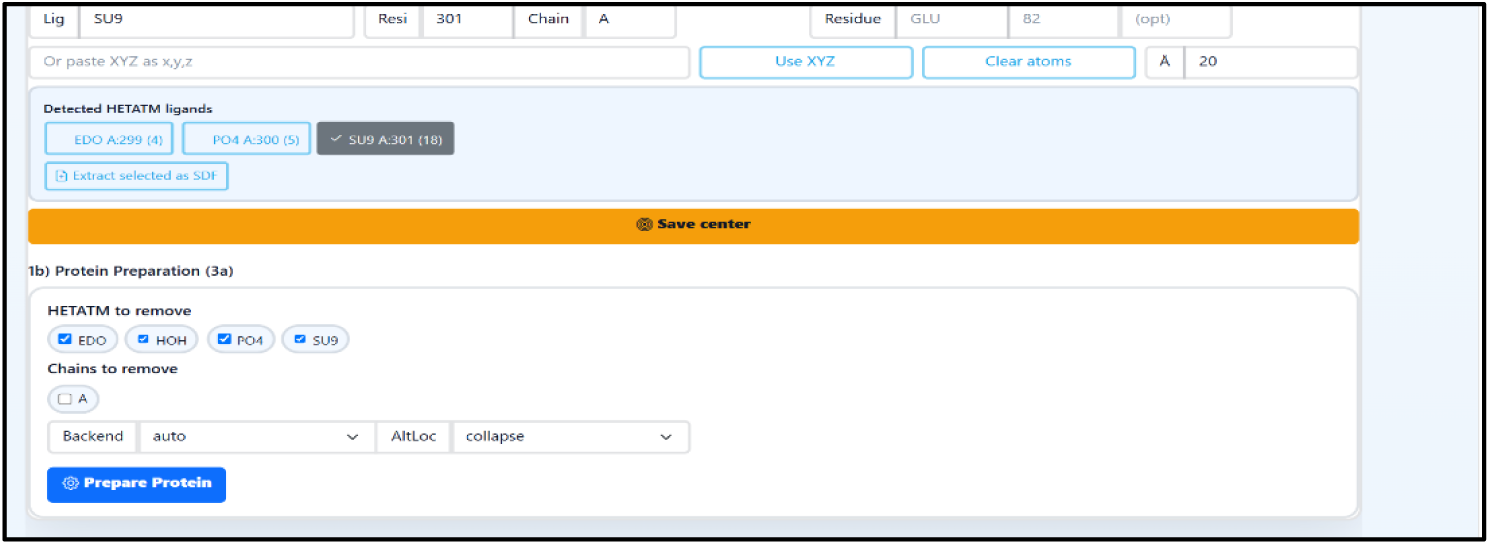
Ligand selection for protein preparation

### Ligand Preparation

Each of the chosen FDA-approved substances from the ZINC20 database was uploaded separately in SDF format to the AutoDock-Vina PrepServer. The compounds were processed for further docking calculations using the ligand-preparation module.

The docking process was set up to produce nine docking poses for each ligand, and 64 ligand conformations were produced for every docking computation. The Auto Dock Vina docking engine was then used to dock the produced ligand into the designated CDK2 binding site. The docking calculation produced several potential ligand-binding orientations and the Docking scores for each molecule. The docking findings were supplied as output files in a ZIP package that can be downloaded. In order to compare and rank the screened compounds, the docking scores and associated ligand data were then recorded.

**Figure 9:**
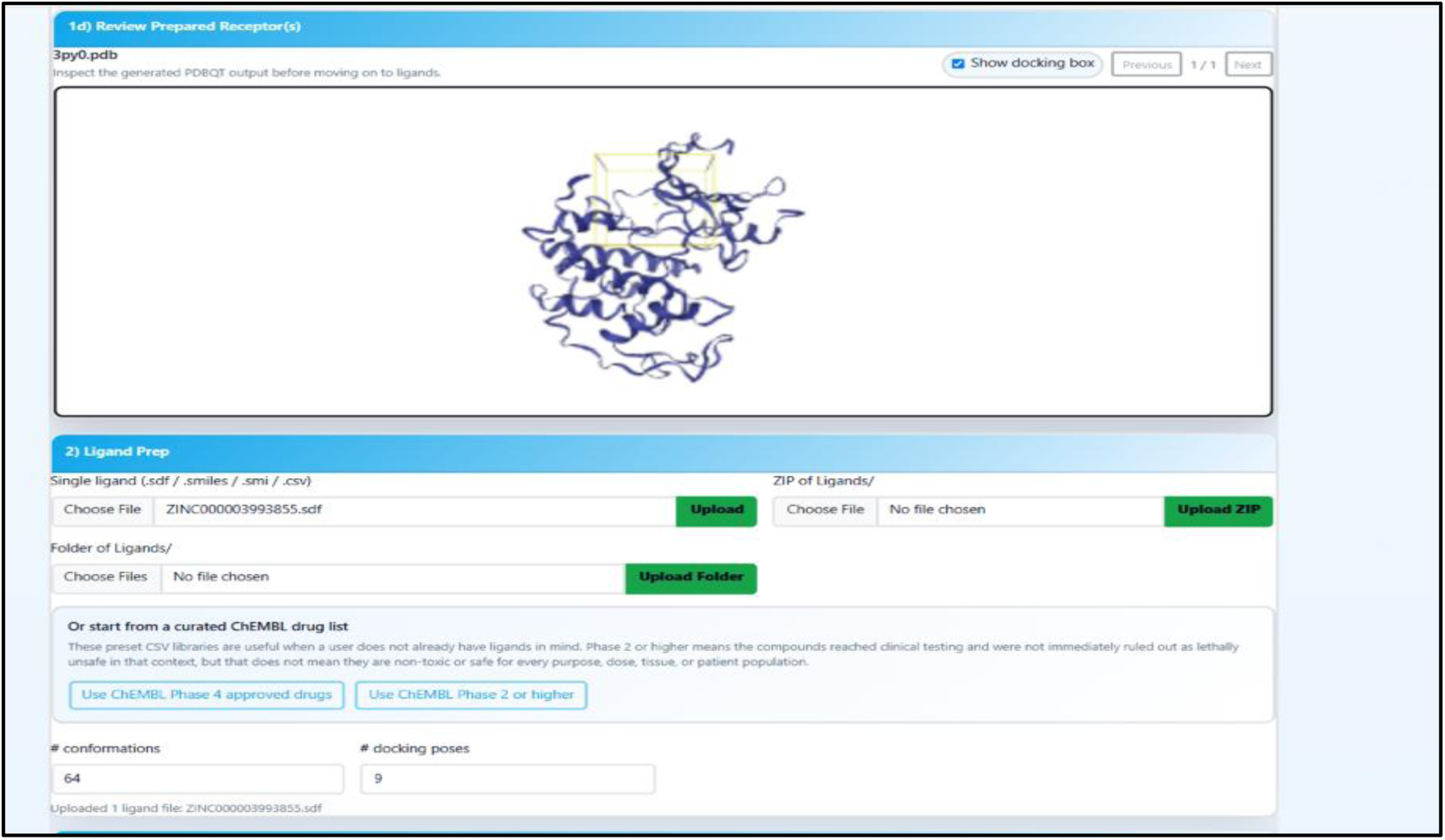
Auto Dock-Vina PrepServer Protein Preparation / Box Generation

### Molecular Docking Using Webina

Molecular docking in Webina(https://durrantlab.pitt.edu/webina/) was then performed using the prepared CDK2 receptor and ligand files produced via the AutoDock-Vina PrepServer methodology. The downloadable ZIP packages created throughout the preparation phase contained the necessary docking files

**Figure 10:**
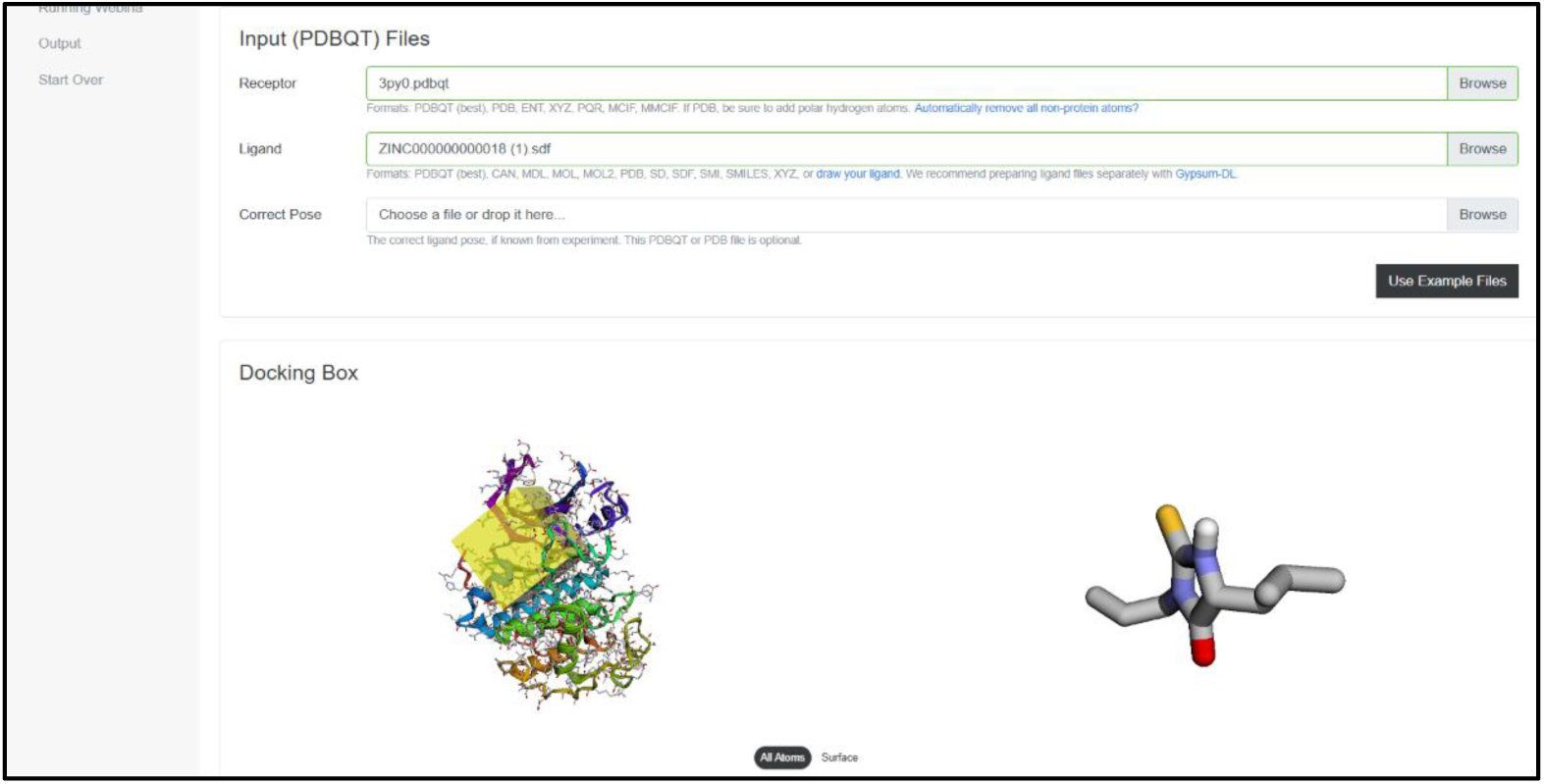
Ligand-binding site of CDK2 in the crystal structure 3PY0

The experimentally identified ligand-binding site of CDK2 in the crystal structure 3PY0 served as the basis for defining the docking box. The ligand-binding area was located using the co-crystallized inhibitor SU9516 (SU9) found in the original crystal structure as a structural reference. The binding pocket that SU9516 inhabited was enclosed by the docking-box centre. The docking-box centre was set at X = −92.14, Y = 79.54, and Z = 85.22 Å. Based on this binding-site a box size of 20 × 20 × 20 Å was used.

**Figure 11:**
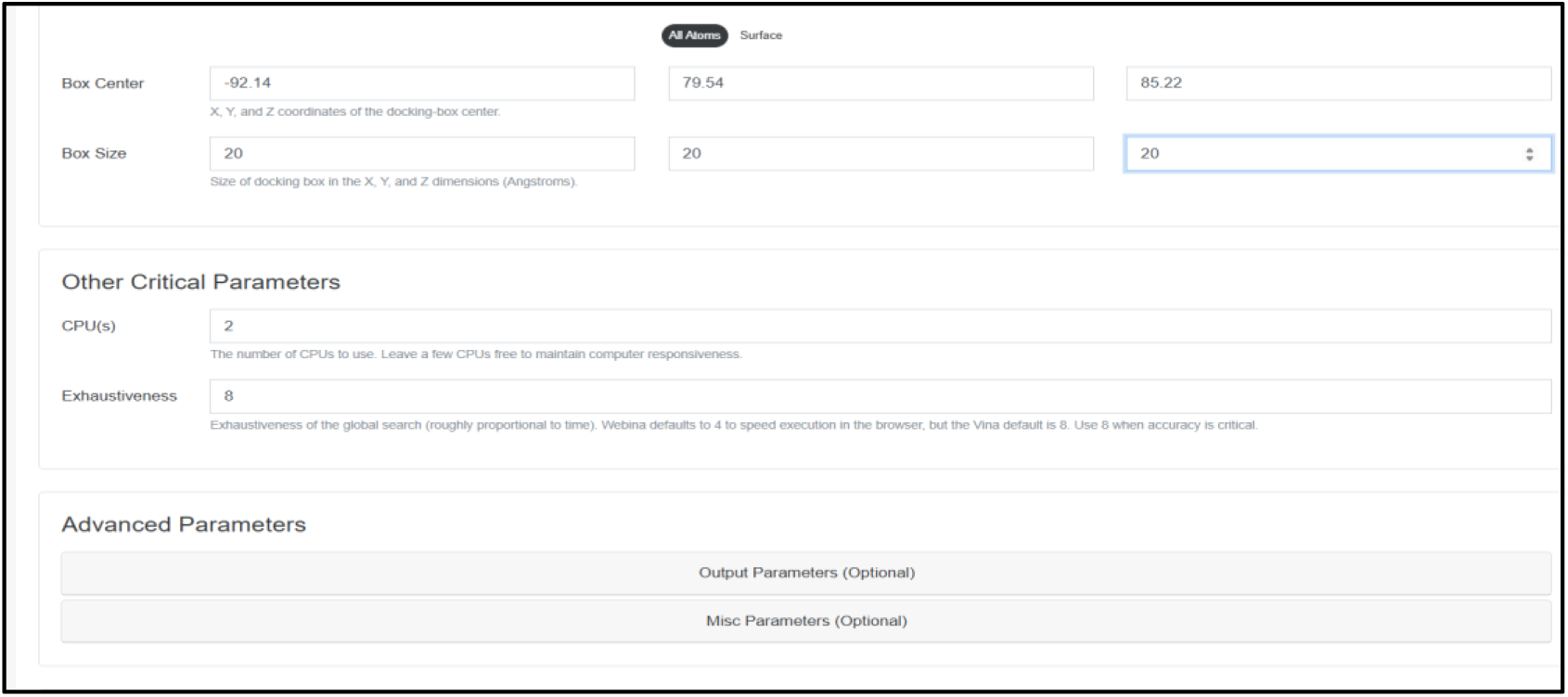
Docking-box centre and critical parameters

Two CPU cores with an exhaustiveness value of eight were used for the Webina docking computation. The projected binding affinity, reported in kcal/mol used to evaluate the docking results. More negative affinity values were thought to indicate higher expected binding. Using PyMOL(https://pymol.org/), PLIP(https://plip-tool.biotec.tu-dresden.de/plip-web/plip/index), the obtained docking poses were used for structural visualization and protein–ligand interaction. The same molecular docking procedure was applied consistently to all selected FDA-approved drugs to ensure comparability of the predicted binding affinities.

### Webina Output Files

The Webina output files with the docked ligand pose and predicted binding-affinity values were downloaded. While the output PDBQT file included. These downloaded files were then utilized for PLIP-based protein–ligand interaction research, PyMOL visualization, and binding-affinity comparison.

### Visualization and Structural Analysis Using PyMOL

PyMOL version 3.1.8 was used for molecular visualization and structural examination of the docked protein–ligand structure from Webina. Before the docked ligand was visible, the generated CDK2 structure was cleaned by eliminating the residual crystallographic elements and solvent molecules. The top-ranked docking pose produced by Webina are used. A stick representation of the docked ligand was used, and a protein–ligand complex was created to show the ligand’s spatial orientation in relation to the receptor. After the protein–ligand complex was created in PyMOL, the complex structure was stored and downloaded in PDB (.pdb) format for later molecular visualization, structural analysis, and PLIP protein–ligand interaction study.

### Protein–Ligand Interaction Analysis Using PLIP

Protein–ligand interactions of the selected docked drug–CDK2 complexes were analysed using the Protein–Ligand Interaction Profiler (PLIP). The PDB-format protein–ligand complex files generated using PyMOL were uploaded to PLIP. Which gives information about hydrogen bonds, hydrophobic interactions, salt bridges, π-stacking, π-cation interactions, and halogen bonds. These were compared with docking scores

## ADMET Filtering

Molecular docking and protein–ligand interaction research once completed ADMET profiling was carried out utilizing the SwissADME(https://swissadme.ch/index.php) and ProTox-3 web servers to further assess the therapeutic candidates. Drug-likeness and pharmacokinetic characteristics were evaluated using SwissADME, with a focus on projected gastrointestinal (GI) absorption and adherence to Lipinski’s Rule of Five. Among the chosen candidates, the acquired physicochemical and pharmacokinetic properties were noted. The ProTox-3 web

server(https://tox.charite.de/protox3/index) was used to assess the toxicity profiles of the chosen candidates. Acute toxicity and possible organ toxicity were projected, along with the relevant toxicity and predicted LD²₀ values. ADMET results were taken into consideration with the docking affinity and protein–ligand interaction patterns.

**Table 3:** 21 screened FDA approved drugs with ZINC ID and docking score

| **S. No.** | **Drug** | **ZINC ID** | **Best docking score (kcal/mol)** |
|---|---|---|---|
| 1 | Baclofen | ZINC000000000061 | −6.813 |
| 2 | Dexbrompheniramine | ZINC000000000096 | −8.443 |
| 3 | Carbinoxamine | ZINC000000000122 | −8.204 |
| 4 | Carteolol | ZINC000000000128 | −8.650 |
| 5 | Formoterol | ZINC856 | −8.865 |
| 6 | Alprazolam | ZINC903 | −5.844 |
| 7 | Apomorphine | ZINC9073 | −11.536 |
| 8 | Exjade | ZINC1481815 | −9.630 |
| 9 | Methotrexate | ZINC1529323 | −9.003 |
| 10 | Lapatinib | ZINC1550477 | −11.273 |
| 11 | Nilotinib | ZINC6716957 | −11.398 |
| 12 | Tadalafil | ZINC3993855 | −9.374 |
| 13 | Cyclopentolate | ZINC196 | −8.159 |
| 14 | Diethylpropion | ZINC215 | −7.168 |
| 15 | Esmolol | ZINC257 | −7.440 |
| 16 | Fenoprofen | ZINC301 | −8.918 |
| 17 | Flurbiprofen | ZINC323 | −9.186 |
| 18 | Cuvposa | ZINC346 | −7.892 |
| 19 | Granisetron | ZINC347 | −8.611 |
| 20 | Labetalol | ZINC416 | −9.290 |
| 21 | Terbutaline | ZINC746 | −7.345 |

**Table 4:** Screened FDA approved drugs with PLIP details with Hydrogen bonds and Hydrophobic interactions with other interaction

| **S. No.** | **Drug** | **Docking score (kcal/mol)** | **Hydrogen bonds** | **Hydrophobic interactions** |
|---|---|---|---|---|
| 1 | Baclofen | −6.813 | 4 | 1 |
| 2 | Dexbrompheniramine | −8.443 | 1 | 7 |
| 3 | Carbinoxamine | −8.204 | 1 | 7 |
| 4 | Carteolol | −8.650 | 3 | 8 |
| 5 | Formoterol | −8.865 | 3 | 7 |
| 6 | Alprazolam | −5.844 | 0 | 5 |
| 7 | Apomorphine | −11.536 | 3 | 8 |
| 8 | Exjade | −9.630 | 2 | 11 |
| 9 | Methotrexate | −9.003 | 4 | 7 |
| 10 | Lapatinib | −11.273 | 3 | 9 |
| 11 | Nilotinib | −11.398 | 1 | 10 |
| 12 | Tadalafil | −9.374 | 5 | 2 |
| 13 | Cyclopentolate | −8.159 | 1 | 8 |
| 14 | Diethylpropion | −7.168 | 0 | 10 |
| 15 | Esmolol | −7.440 | 1 | 10 |
| 16 | Fenoprofen | −8.918 | 1 | 10 |
| 17 | Flurbiprofen | −9.186 | 2 | 8 |
| 18 | Cuvposa | −7.892 | 0 | 12 |
| 19 | Granisetron | −8.611 | 0 | 8 |
| 20 | Labetalol | −9.290 | 4 | 10 |
| 21 | Terbutaline | −7.345 | 4 | 9 |

**Table 5:**
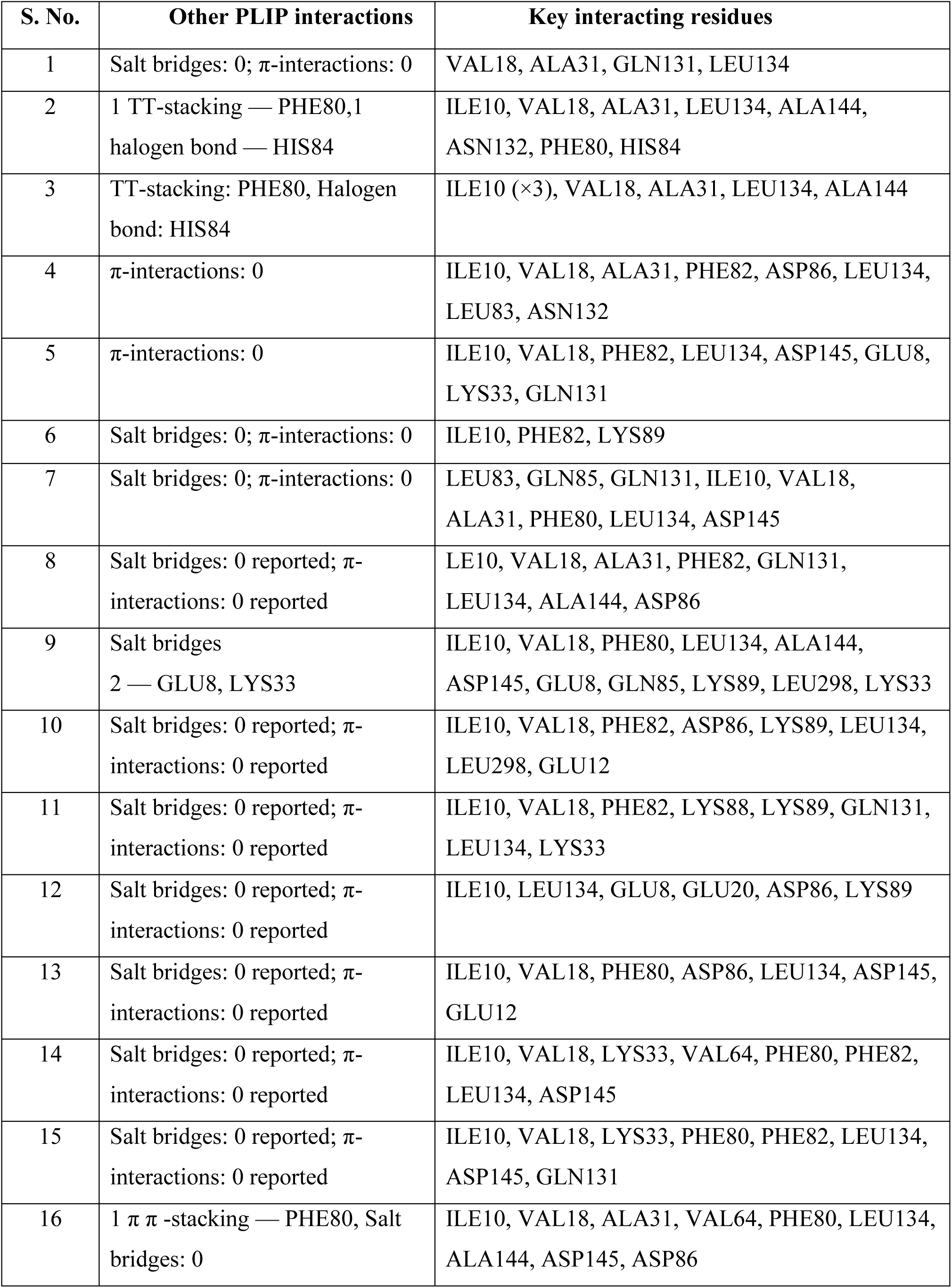
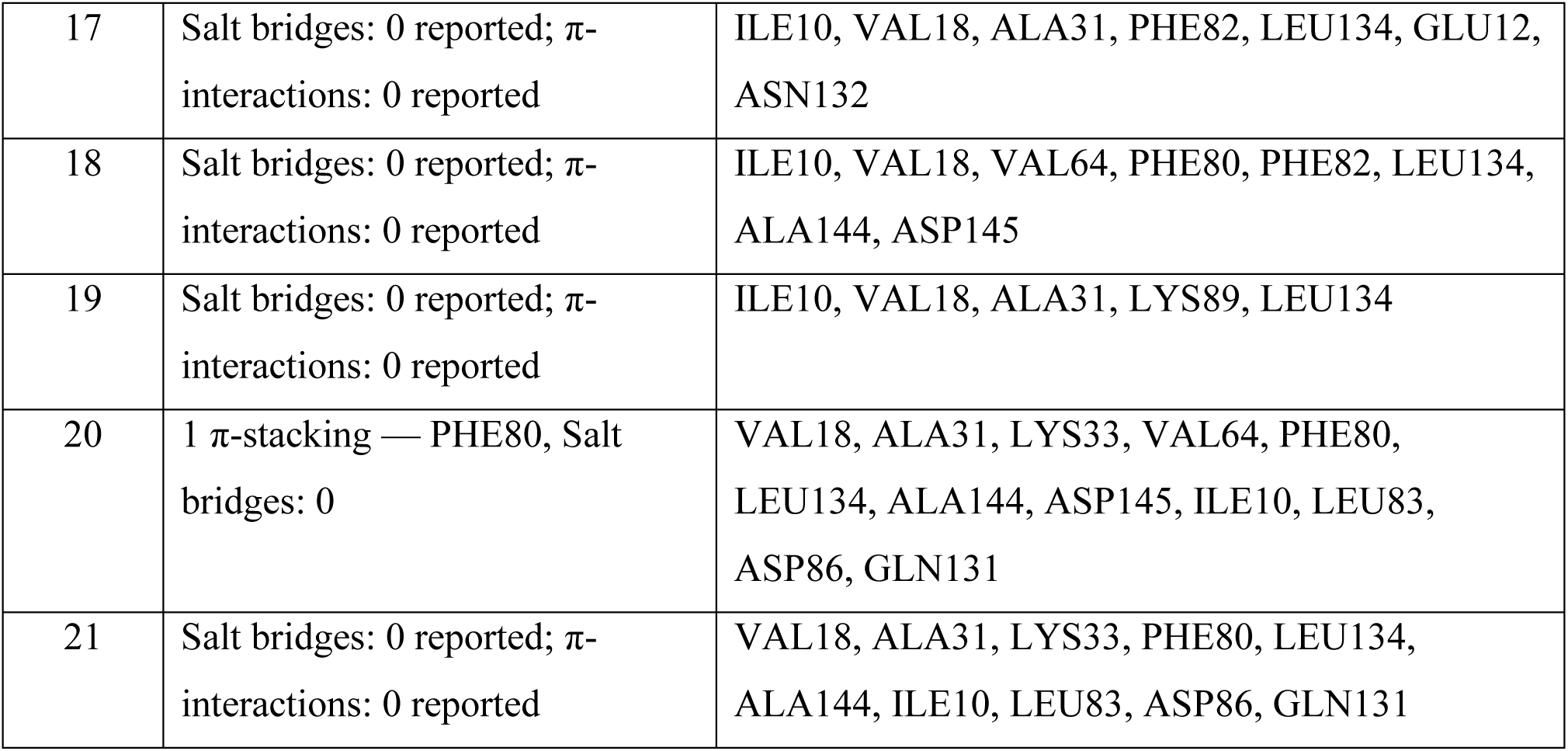
Screened FDA approved drugs with Key interacting residues

| S. No. | Other PLIP interactions | Key interacting residues |
|---|---|---|
| 1 | Salt bridges: 0; π-interactions: 0 | VAL18, ALA31, GLN131, LEU134 |
| 2 | 1 TT-stacking — PHE80,1 halogen bond — HIS84 | ILE10, VAL18, ALA31, LEU134, ALA144, ASN132, PHE80, HIS84 |
| 3 | TT-stacking: PHE80, Halogen bond: HIS84 | ILE10 (×3), VAL18, ALA31, LEU134, ALA144 |
| 4 | π-interactions: 0 | ILE10, VAL18, ALA31, PHE82, ASP86, LEU134, LEU83, ASN132 |
| 5 | π-interactions: 0 | ILE10, VAL18, PHE82, LEU134, ASP145, GLU8, LYS33, GLN131 |
| 6 | Salt bridges: 0; π-interactions: 0 | ILE10, PHE82, LYS89 |
| 7 | Salt bridges: 0; π-interactions: 0 | LEU83, GLN85, GLN131, ILE10, VAL18, ALA31, PHE80, LEU134, ASP145 |
| 8 | Salt bridges: 0 reported; π-interactions: 0 reported | LE10, VAL18, ALA31, PHE82, GLN131, LEU134, ALA144, ASP86 |
| 9 | Salt bridges<br>2 — GLU8, LYS33 | ILE10, VAL18, PHE80, LEU134, ALA144, ASP145, GLU8, GLN85, LYS89, LEU298, LYS33 |
| 10 | Salt bridges: 0 reported; π-interactions: 0 reported | ILE10, VAL18, PHE82, ASP86, LYS89, LEU134, LEU298, GLU12 |
| 11 | Salt bridges: 0 reported; π-interactions: 0 reported | ILE10, VAL18, PHE82, LYS88, LYS89, GLN131, LEU134, LYS33 |
| 12 | Salt bridges: 0 reported; π-interactions: 0 reported | ILE10, LEU134, GLU8, GLU20, ASP86, LYS89 |
| 13 | Salt bridges: 0 reported; π-interactions: 0 reported | ILE10, VAL18, PHE80, ASP86, LEU134, ASP145, GLU12 |
| 14 | Salt bridges: 0 reported; π-interactions: 0 reported | ILE10, VAL18, LYS33, VAL64, PHE80, PHE82, LEU134, ASP145 |
| 15 | Salt bridges: 0 reported; π-interactions: 0 reported | ILE10, VAL18, LYS33, PHE80, PHE82, LEU134, ASP145, GLN131 |
| 16 | 1 π π -stacking — PHE80, Salt bridges: 0 | ILE10, VAL18, ALA31, VAL64, PHE80, LEU134, ALA144, ASP145, ASP86 |
| 17 | Salt bridges: 0 reported; π-interactions: 0 reported | ILE10, VAL18, ALA31, PHE82, LEU134, GLU12, ASN132 |
| 18 | Salt bridges: 0 reported; π-interactions: 0 reported | ILE10, VAL18, VAL64, PHE80, PHE82, LEU134, ALA144, ASP145 |
| 19 | Salt bridges: 0 reported; π-interactions: 0 reported | ILE10, VAL18, ALA31, LYS89, LEU134 |
| 20 | 1 π-stacking — PHE80, Salt bridges: 0 | VAL18, ALA31, LYS33, VAL64, PHE80, LEU134, ALA144, ASP145, ILE10, LEU83, ASP86, GLN131 |
| 21 | Salt bridges: 0 reported; π-interactions: 0 reported | VAL18, ALA31, LYS33, PHE80, LEU134, ALA144, ILE10, LEU83, ASP86, GLN131 |

**Table 6:** Swiss ADME result

| Ligand | MW (g/mol) | TPSA (Å<sup>2</sup>) | GI absorption | BBB permeant | P-gp substrate | Lipinski violations | Bioavailability score | PAINS alerts | Brenk alerts | Synthetic accessibility |
|---|---|---|---|---|---|---|---|---|---|---|
| Apomorphine | 267.32 | 43.7 | High | Yes | Yes | 0 | 0.55 | 1 | 1 | 3.26 |
| Baclofen | 213.66 | 63.32 | High | Yes | No | 0 | 0.55 | 0 | 0 | 1.81 |
| Dexbrompheniramine | 319.24 | 16.13 | High | Yes | No | 0 | 0.55 | 0 | 0 | 2.77 |
| Carbinoxamine | 290.79 | 25.36 | High | Yes | No | 0 | 0.55 | 0 | 0 | 2.98 |
| Carteolol | 292.37 | 70.59 | High | No | Yes | 0 | 0.55 | 0 | 0 | 3.12 |
| Formoterol | 344.4 | 97.5 | High | No | Yes | 0 | 0.55 | 0 | 2 | 3.08 |
| Alprazolam | 308.76 | 43.07 | High | Yes | Yes | 0 | 0.55 | 0 | 0 | 3.33 |
| Exjade | 373.36 | 108.47 | High | No | No | 0 | 0.56 | 0 | 0 | 2.95 |
| Methotrexate | 454.44 | 210.54 | Low | No | No | 1 | 0.11 | 0 | 0 | 3.58 |
| Lapatinib | 581.06 | 114.73 | Low | No | No | 1 | 0.55 | 0 | 0 | 4.05 |
| Nilotinib | 529.52 | 97.62 | Low | No | Yes | 1 | 0.55 | 0 | 0 | 3.81 |
| Tadalafil | 389.4 | 74.87 | High | No | No | 0 | 0.55 | 1 | 0 | 3.82 |
| Cyclopentolate | 291.39 | 49.77 | High | Yes | No | 0 | 0.55 | 0 | 0 | 3.19 |
| Diethylpropion | 205.3 | 20.31 | High | Yes | No | 0 | 0.55 | 0 | 0 | 1.78 |
| Esmolol | 295.37 | 67.79 | High | Yes | No | 0 | 0.55 | 0 | 0 | 2.68 |
| Fenoprofen | 242.27 | 46.53 | High | Yes | No | 0 | 0.85 | 0 | 0 | 2.41 |
| Flurbiprofen | 244.26 | 37.3 | High | Yes | No | 0 | 0.85 | 0 | 0 | 2.45 |
| Cuvposa | 318.43 | 46.53 | High | Yes | Yes | 0 | 0.55 | 0 | 1 | 3.7 |
| Granisetron | 312.41 | 50.16 | High | Yes | Yes | 0 | 0.55 | 0 | 0 | 4.08 |
| Labetalol | 328.41 | 95.58 | High | No | Yes | 0 | 0.55 | 0 | 0 | 3 |
| Terbutaline | 225.28 | 72.72 | High | No | No | 0 | 0.55 | 0 | 0 | 2.13 |

**Table 7:** Results showing toxicity classes

| **Ligand** | **LD50(mg/kg)** | **Toxicity Class** | **H** | **C** | **M** | **Cy** | **I** |
|---|---|---|---|---|---|---|---|
| Baclofen | 145 | 3 | - | + | - | - | - |
| Dexbrompheniramine | 147 | 3 | - | - | - | - | + |
| Carbinoxamine | 162 | 3 | - | - | - | - | + |
| Carteolol | 810 | 4 | - | - | - | - | + |
| Formoterol | 3130 | 5 | - | - | - | - | + |
| Alprazolam | 770 | 4 | - | - | - | - | - |
| Apomorphine | 100 | 3 | - | - | + | - | + |
| Exjade | 1000 | 4 | + | + | - | - | - |
| Methotrexate | 3 | 1 | + | - | - | - | - |
| Lapatinib | 1500 | 4 | + | - | - | + | + |
| Nilotinib | 800 | 4 | + | + | - | - | + |
| Tadalafil | 300 | 3 | - | + | - | + | + |
| Cyclopentolate | 960 | 4 | - | - | - | - | + |
| Diethylpropion | 160 | 3 | - | - | - | - | - |
| Esmolol | 2000 | 4 | - | - | - | - | - |
| Fenoprofen | 800 | 4 | + | - | - | - | - |
| Flurbiprofen | 117 | 3 | + | + | - | - | - |
| Cuvposa | 500 | 4 | - | - | - | - | - |
| Granisetron | 350 | 4 | - | + | + | - | - |
| Labetalol | 660 | 4 | + | - | - | - | - |
| Terbutaline | 205 | 3 | - | - | - | - | - |

## Results

### Molecular Docking and Protein–Ligand Interaction Analysis of Top-Ranked Candidates

Following the initial screening of FDA-approved drugs compounds were selected based on their predicted toxicity class and predefined drug-likeness and ADME criteria obtained from Swiss ADME and ProTox. Most Promising class 4 Compounds that satisfied the major selection criteria were prioritized, while compounds showing certain limitations were retained only when their overall computational profile remained promising. Based on *this Lapatinib, Flurbiprofen, Fenoprofen, Carteolol and Granisetron* were Recognised. The selected candidates were subsequently evaluated more deeply based on their docking scores, predicted binding poses, and interactions identified using PLIP. Out of this top candidate was selected.

**Figure 12:**
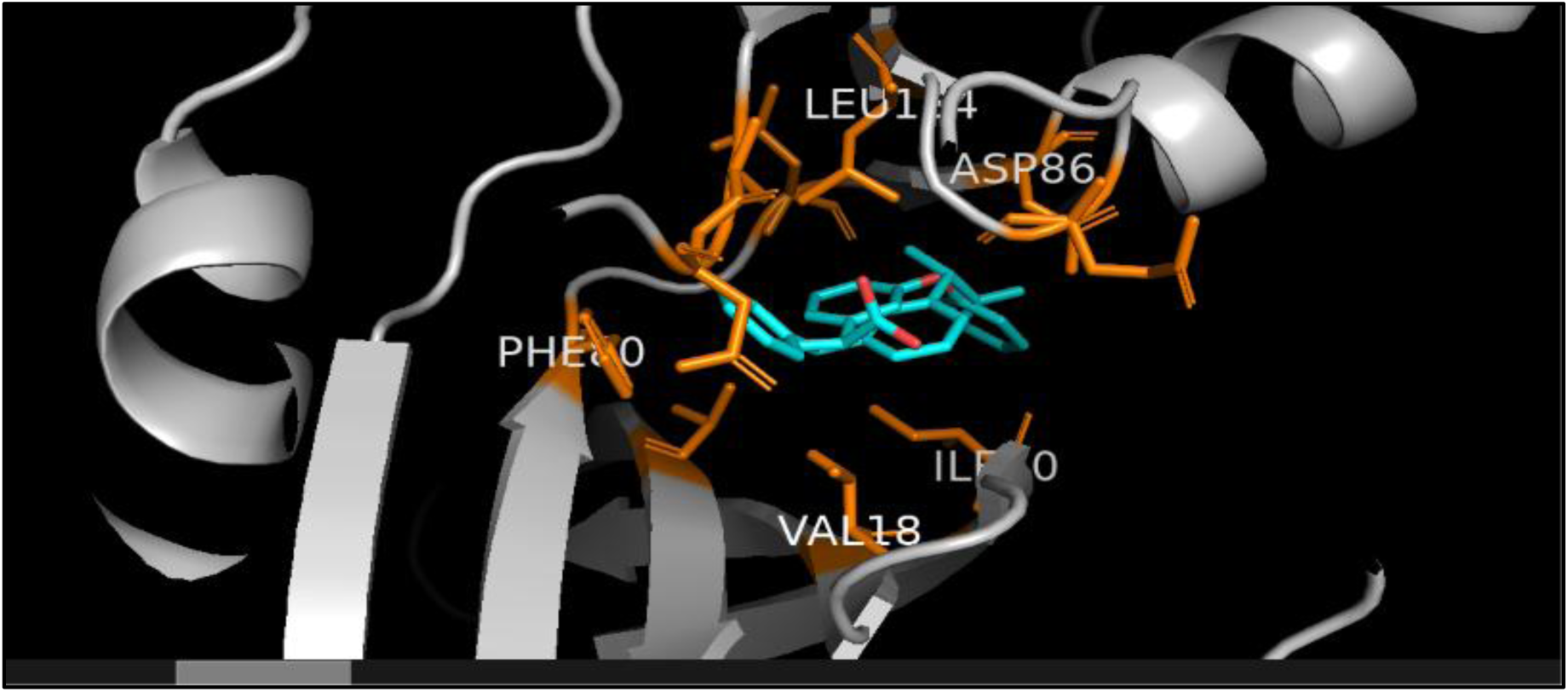
Fenoprofen –CDK2 Molecular Docking and Interaction Analysis (PHE80 → orange; π-stacking with Fenoprofen, ASP86 → orange; hydrogen bond with Fenoprofen, LEU134 → orange; hydrophobic interaction)

**Figure 13:**
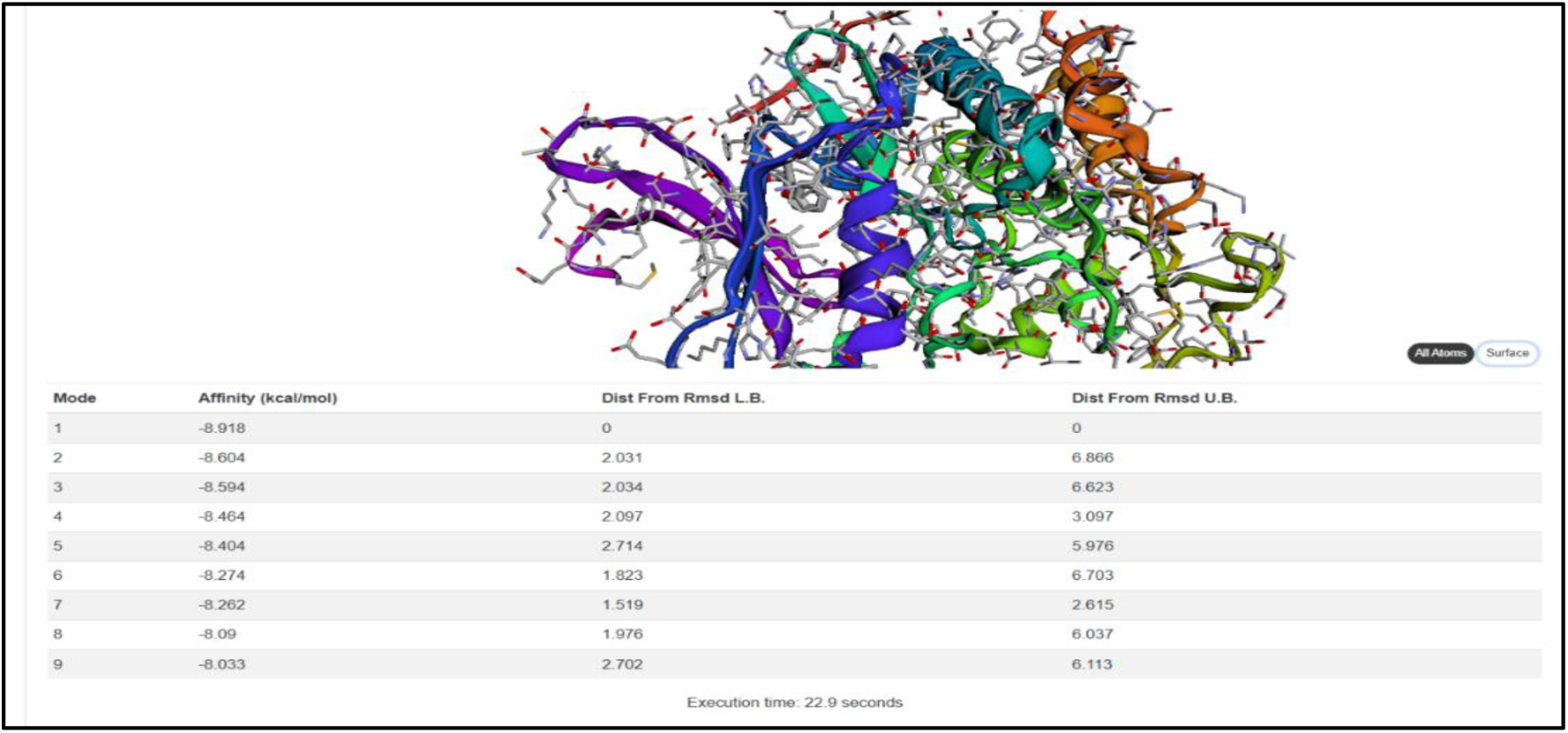
Webina Docking Scores of Fenoprofen –CDK2 Complex (Modes1-9)

The best docking pose (Mode 1) of Fenoprofen showed a docking score of −8.918 kcal/mol against CDK2. PLIP analysis identified hydrophobic interactions with several binding-pocket residues, a hydrogen bond involving ASP86, and a π-stacking interaction with PHE80. However, the interaction profile does not indicate exceptionally strong binding, as only one hydrogen bond and one π-stacking interaction were observed. Therefore, PHE80, ASP86, and LEU134 were selected for visualization as key interacting residues contributing to the predicted binding of Fenoprofen within the CDK2 pocket and BBB permeant is not convincing with Hepatotoxicity active

**Figure 14:**
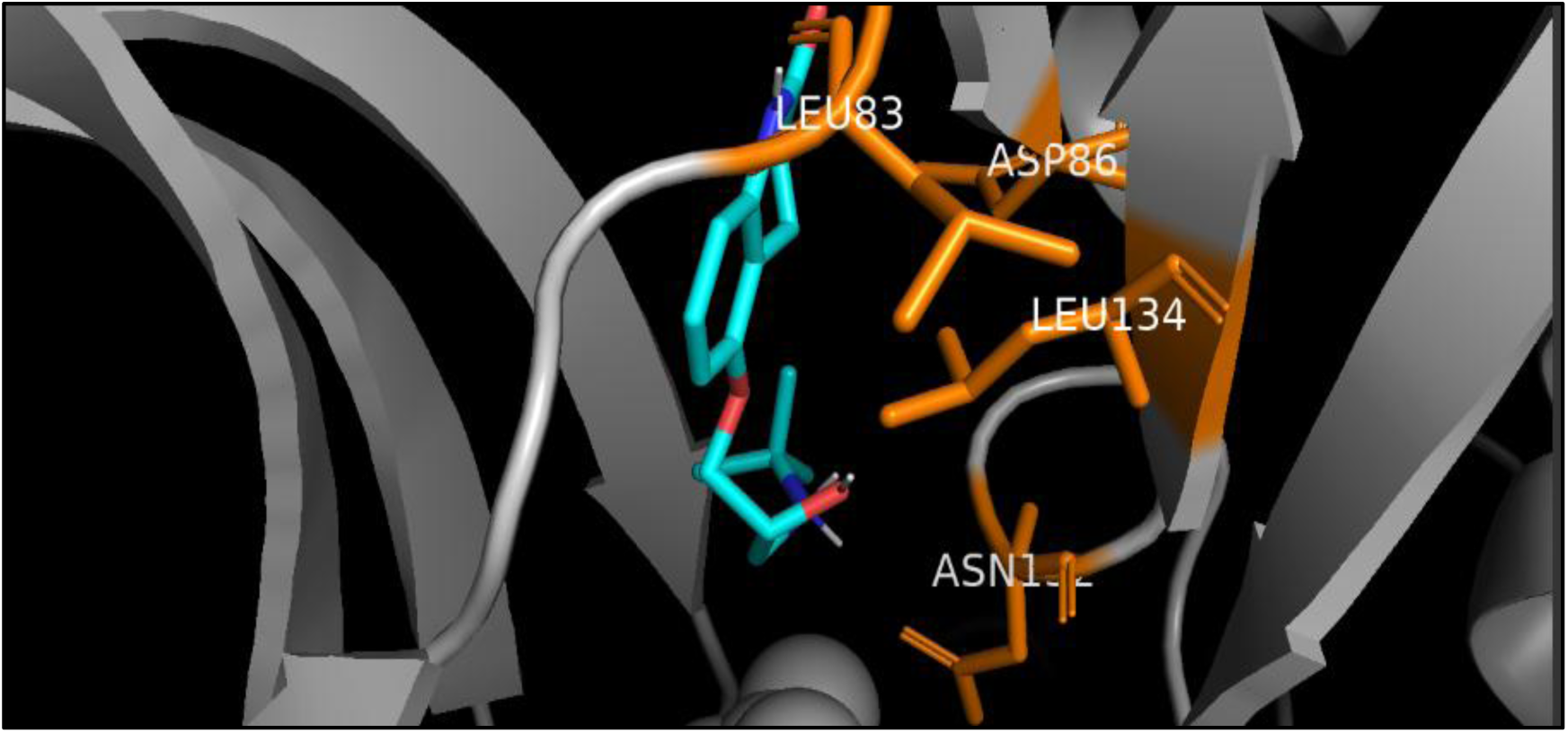
Carteolol-CDK2 Molecular Docking and Interaction Analysis (Orange → key interacting residues: LEU83, ASP86, ASN132, LEU134)

**Figure 15:**
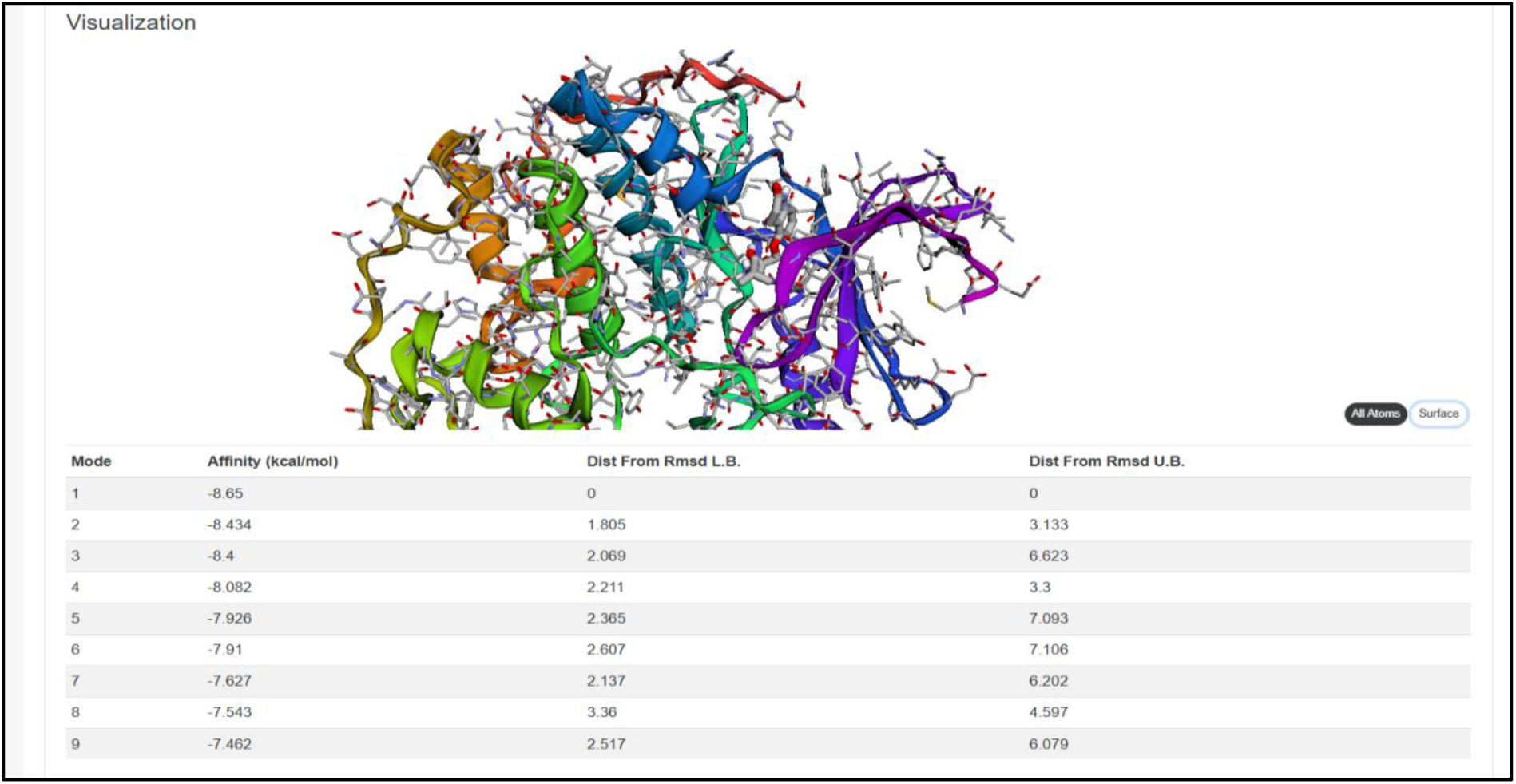
Webina Docking Scores of Carteolol–CDK2 Complex (Modes1-9)

The best docking pose (Mode 1) of Carteolol was visualized in PyMOL to examine its predicted interaction with the CDK2 ATP-binding pocket. PLIP analysis identified hydrogen-bond interactions with LEU83, ASP86, and ASN132, along with hydrophobic interactions involving several pocket residues, including ILE10, VAL18, ALA31, PHE82, ASP86, and LEU134. Therefore, LEU83, ASP86, ASN132, and LEU134 were highlighted as key interacting residues in the 3D structure which provides good structural support for its predicted interaction with CDK2 with good toxicity, all 5 parameters are inactive (carcinogenicity, mutagenicity, cytotoxicity, hepatotoxicity, immunotoxicity) and SWISSadme parameters are also convincing.

**Figure 16:**
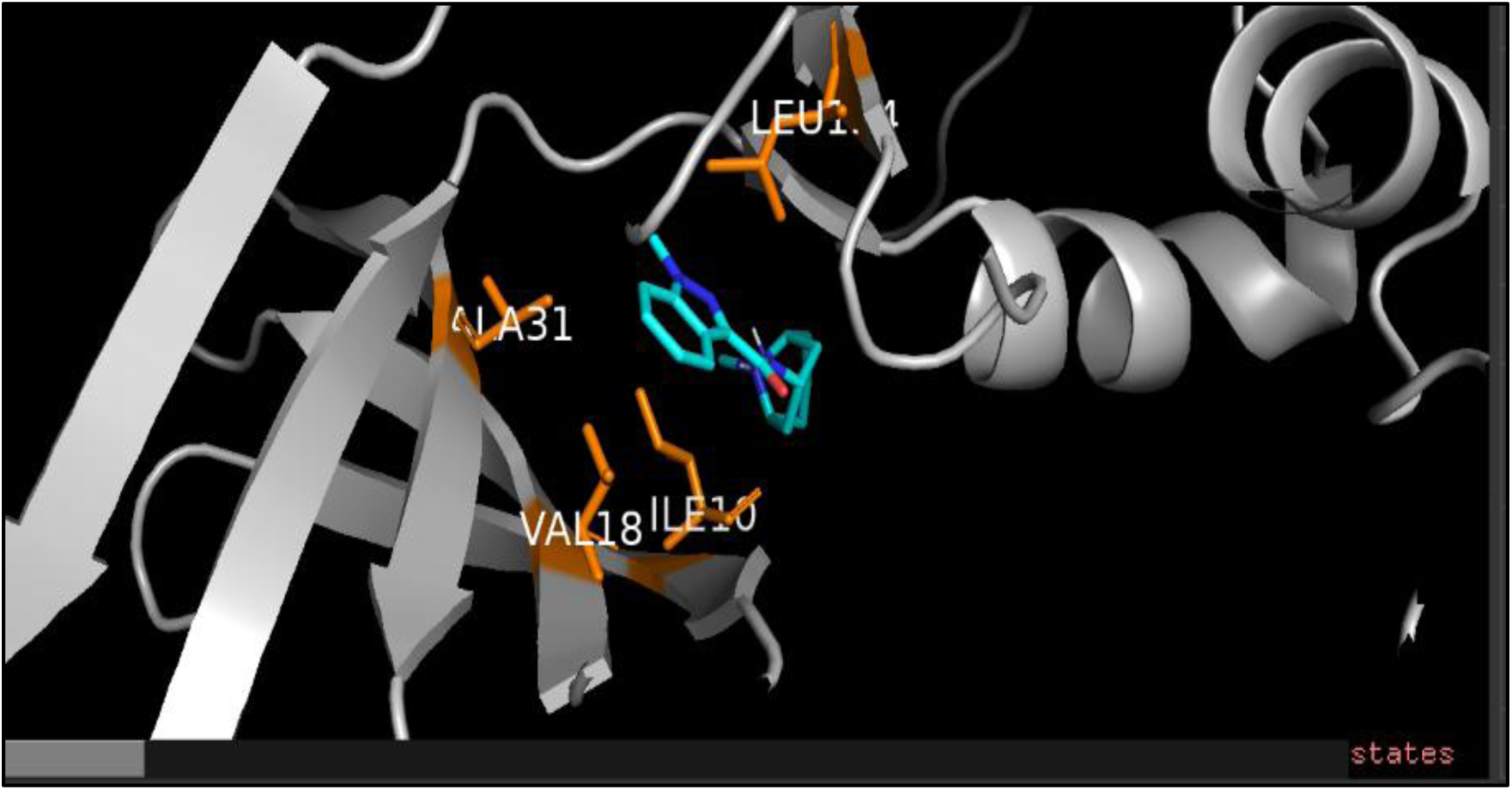
Granisetron –CDK2 Molecular Docking and Interaction Analysis (Orange → key interacting residues: → ILE10, VAL18, ALA31, LEU134)

The best docking pose (Mode 1) of Granisetron showed a favourable docking score of −8.611kcal/mol against CDK2. PLIP analysis identified eight hydrophobic interactions involving ILE10, VAL18, ALA31, LYS89, and LEU134. However, no hydrogen bonds, π-stacking, or salt-bridge interactions were detected in the provided PLIP analysis. Therefore, Granisetron demonstrates a favourable predicted binding pose but a comparatively less specific interaction profile

**Figure 17:**
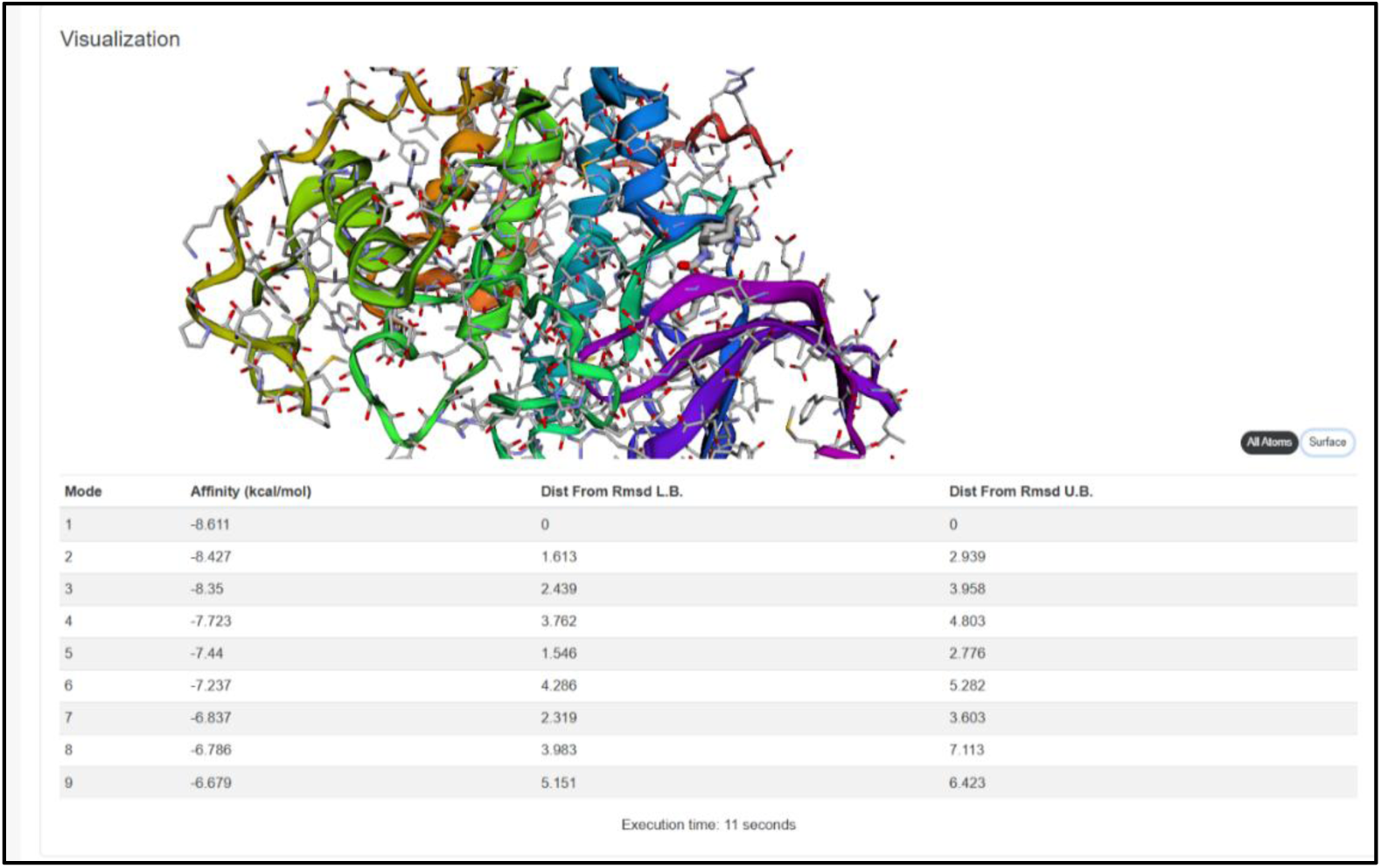
Webina Docking Scores of Granisetron –CDK2 Complex (Modes1-9)

**Figure 18:**
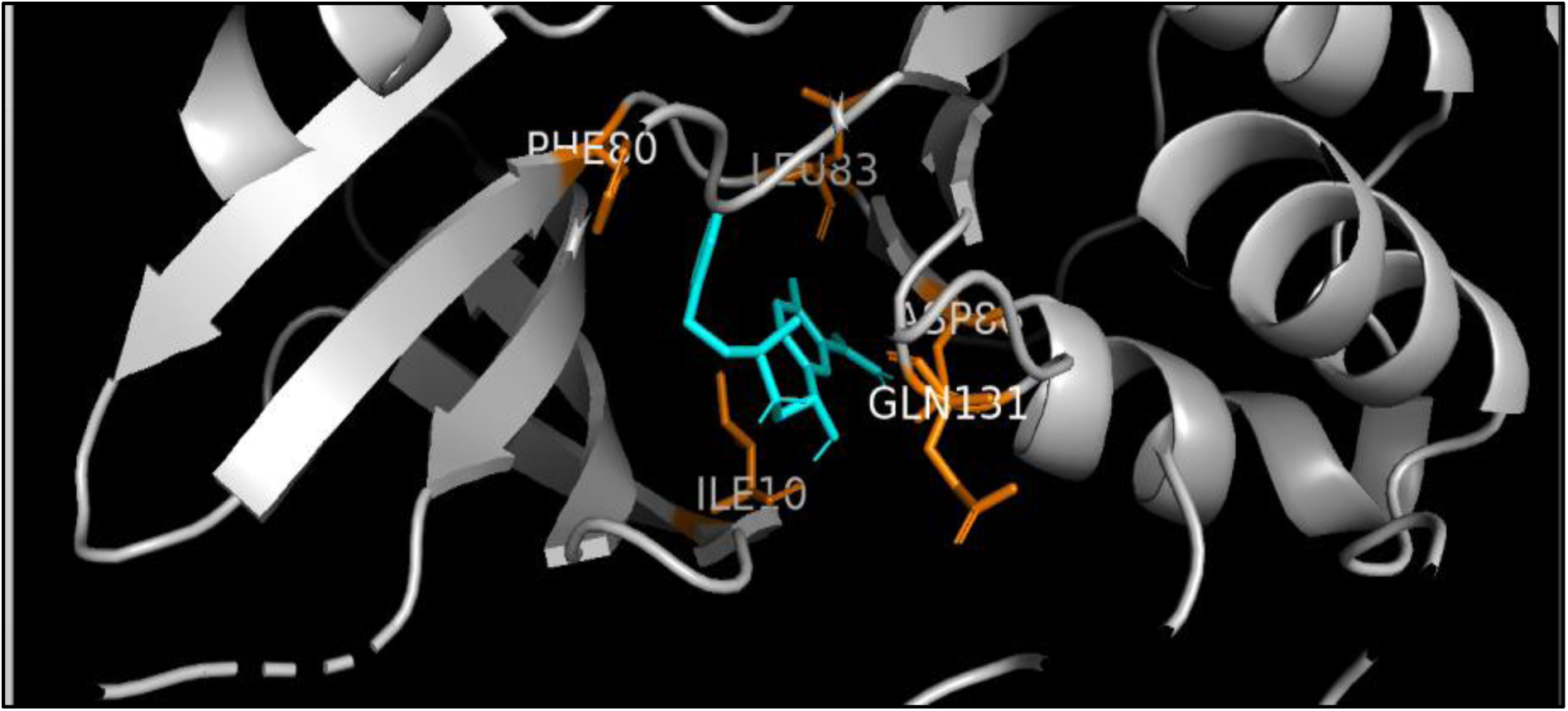
Labetalol–CDK2 Molecular Docking and Interaction Analysis

The best docking pose of Labetalol showed a favourable docking score of −9.290 kcal/mol against CDK2. PLIP analysis identified 10 hydrophobic contacts, four hydrogen bonds involving ILE10, LEU83, ASP86, and GLN131, and a π-stacking interaction with PHE80. These multiple noncovalent interactions support a favourable and well-supported predicted binding mode of Labetalol within the CDK2 binding pocket, indicating its potential as a candidate for further CDK2-targeted drug repurposing studies but has concerns with Swiss ADME and toxicity

**Figure 19:**
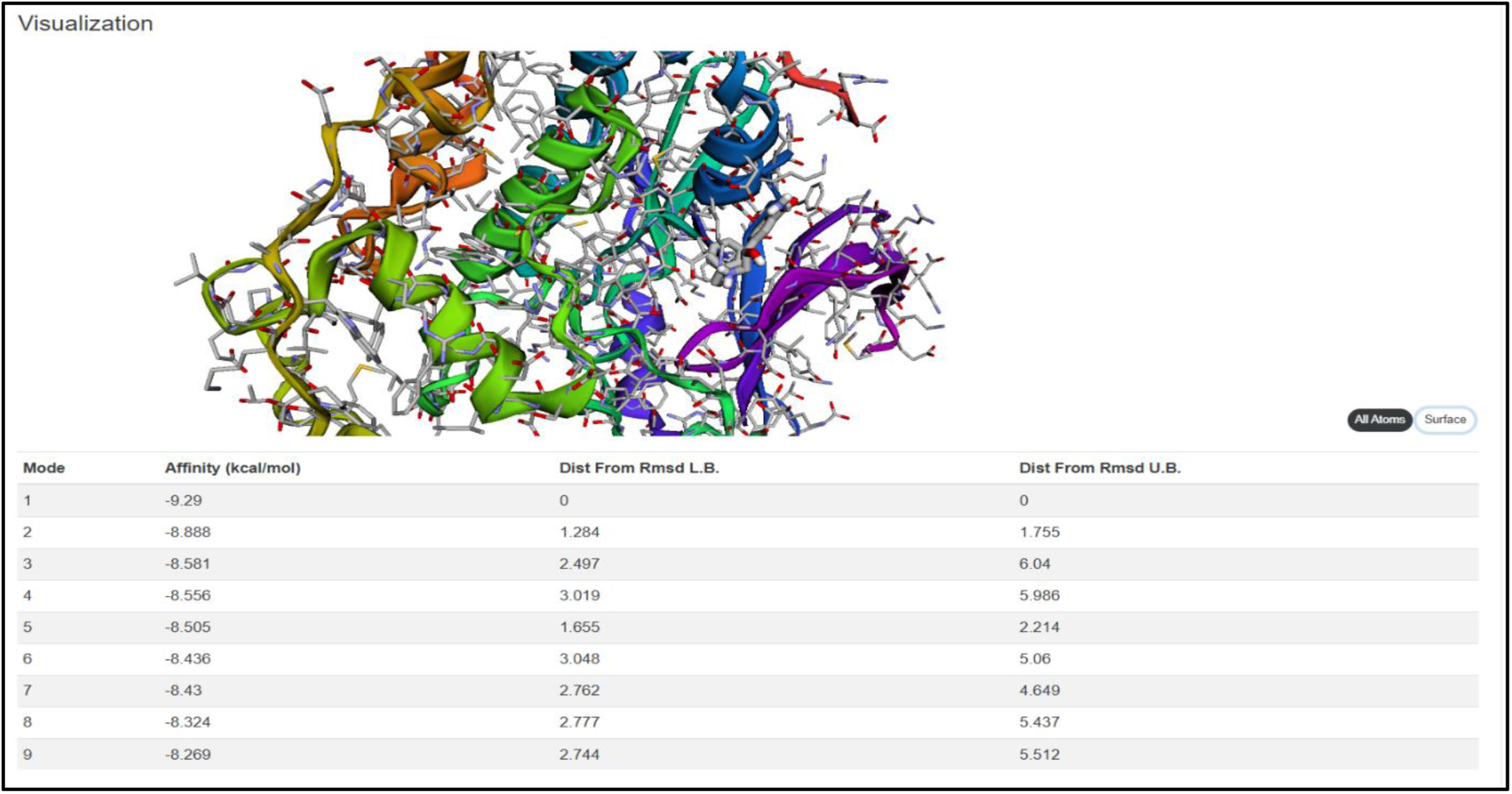
Webina Docking Scores of Labetalol–CDK2 Complex (Modes1-9)

**Figure 20:**
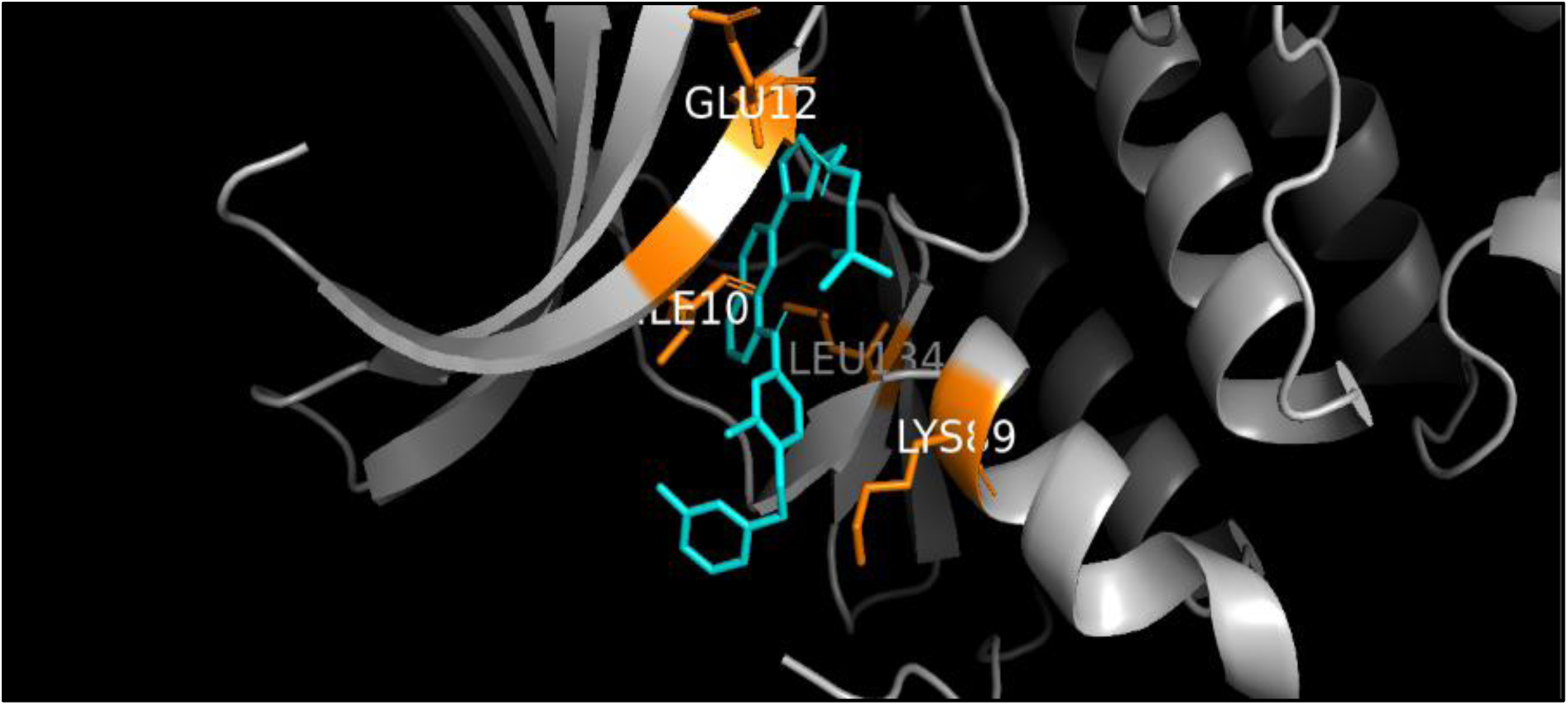
Lapatinib –CDK2 Molecular Docking and Interaction Analysis

**Figure 21:**
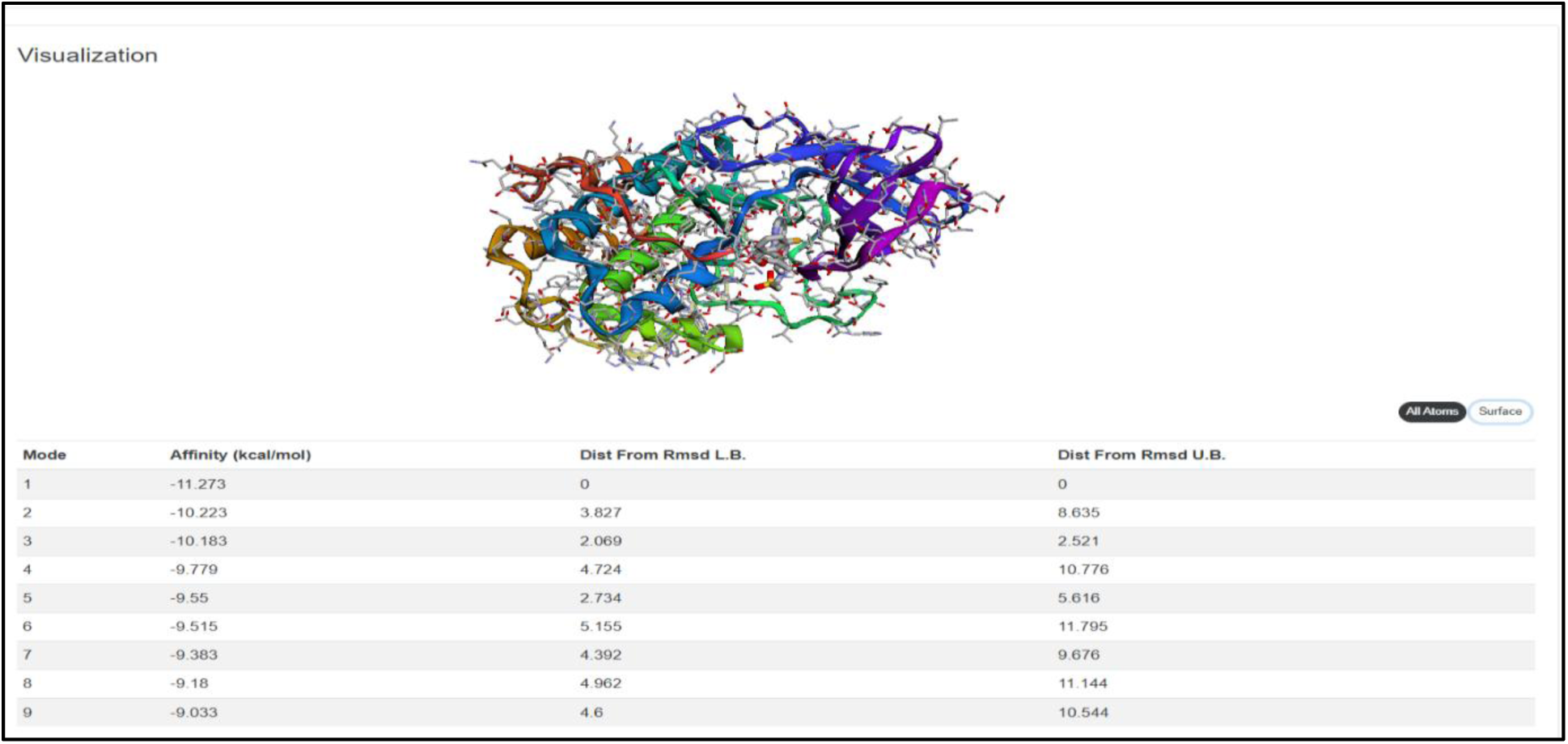
Webina Docking Scores of Lapatinib –CDK2 Complex (Modes1-9)

The best docking pose (Mode 1) of Lapatinib was visualized against CDK2 using PyMOL. PLIP analysis identified nine hydrophobic interactions involving residues ILE10, VAL18, PHE82, ASP86, LYS89, LEU134, and LEU298, along with three hydrogen bonds involving ILE10, GLU12, and LYS89. The docking pose showed multiple interactions within the CDK2 binding pocket, supporting a favourable predicted binding mode of Lapatinib. ILE10, GLU12, LYS89, and LEU134 were highlighted as key interacting residues in the 3D visualization. Overall, the interaction profile suggests that Lapatinib can form a well-supported predicted interaction network within the CDK2 binding pocket.

## Discussion

Study about repurposing of FDA-approved drugs against cyclin-dependent kinase 2 (CDK2) with computational flow includes molecular docking, Protein–Ligand Interaction Profiler (PLIP) analysis, SwissADME and ProTox toxicity prediction. In drug repurposing compounds with best predicted binding score may also have some limitations like toxicity, pharmacokinetics or chemical drug-likeness. Among the screened compounds, Apomorphine produced the strongest docking score at −11.536 kcal/mol, followed by Nilotinib at −11.398 kcal/mol and Lapatinib at −11.273 kcal/mol. These results indicate that several FDA-approved drugs may theoretically fit favourably within the selected CDK2 binding region. Docking scores alone cannot establish whether a compound will function as a biologically active CDK2 inhibitor. PLIP analysis was used to examine the specific interactions and toxicity analysis. Lapatinib was among the strongest docking candidates in the present screening, with a docking score of −11.273 kcal/mol. PLIP analysis identified 12 interactions, consisting of nine hydrophobic interactions and three hydrogen bonds. Lapatinib is an established dual EGFR/HER2 tyrosine-kinase inhibitor. Chu *et al*., 2005 reported that Lapatinib with tamoxifen increased p27 association with cyclin E–CDK2 and inhibition of cyclin E–CDK2 activity in breast cancer models with cell-cycle arrest. This finding is relevant to the present study Lapatinib can influence a biological pathway involving cyclin E–CDK2 but did not provide direct evidence of CDK2 inhibitor. Nilotinib produced the second-strongest docking score overall (−11.398 kcal/mol) in the present study PLIP analysis identified 11 interactions.

Previous experimental research also provides a biological connection between Nilotinib and CDK2-associated cell-cycle regulation. Pham *et al.,* (2024) investigated Nilotinib in human melanoma A375P cells and it can inhibit proliferation and impaired the G1/S transition. *Pham et al.,* did not demonstrate direct inhibition of the CDK2 ATP-binding pocket. Labetalol produced a docking score of −9.290 kcal/mol PLIP analysis identified 15 interactions. These included 10 hydrophobic interactions, four hydrogen bonds and one π-stacking interaction involving PHE80. Phe80 as gatekeeper residue to the hydrophobic environment of the CDK2 ATP-binding. Previous studies also reported interactions involving Phe80 together with other residues such as Ile10, Val18, Leu83 and Leu134 in CDK2 ligand complexes.

SwissADME results supported the drug-likeness of Labetalol zero Lipinski violations, zero PAINS alerts, zero Brenk alerts and high predicted gastrointestinal absorption. Fenoprofen demonstrated a strong and balanced computational profile, with a docking score of −8.918 kcal/mol, 12 PLIP interactions, favourable drug-likeness and a predicted LD50 of 800 mg/kg. Nevertheless, several toxicity endpoints were predicted to be active. Su *et al.,* (2020) reported that Carteolol induced G1 cell-cycle arrest and apoptosis in human corneal endothelial cells.

Carteolol demonstrated a docking score of −8.650 kcal/mol and 11 PLIP interactions, comprising eight hydrophobic interactions and three hydrogen bonds. Include residues LEU83, ASP86 and ASN132 Favours CDK2 binding region. protein–ligand interactions with high predicted gastrointestinal absorption, zero Lipinski violations, absence of PAINS and Brenk alerts, and a comparatively favourable predicted toxicity profile. Its LD50 of 810 mg/kg (Class 4) is considered best among all. Remaining screened drugs were not in consideration which may include reasons of favourable docking scores or interaction profiles, predicted toxicity.

From above details class 4 toxicity group was considered for detailed analysis where Alprazolam has a PAINS alert, In Exjade it was found that hepatotoxicity and carcinogenicity are active. Whereas in Lapatinib (−11.273), Nilotinib (−11.398) has a strong docking score but has low GI absorption level with toxicity concerns. Cyclopentolate and Esmolol Swiss ADME are not in range, Fenoprofen has BBB alert and Cuvposa have Brenk alert 1, Granisetron with toxicity alerts. Out of these Carteolol surpasses toxicity class and predefined drug-likeness and ADME criteria obtained from SwissADME and ProTox and shows favourable predicted binding pose. The present computational findings suggest that **Carteolol** may have potential for Repurposing as a CDK2-targeting compound. Its predicted binding within the CDK2 pocket, together with multiple hydrogen-bond and hydrophobic interactions, supports further investigation of its potential CDK2 inhibitory activity

### Conclusion and Future scope

FDA-approved drugs for their potential repurposing against cyclin-dependent kinase 2 (CDK2) using computational work flow says results demonstrated not on based on docking scores but also rely on pharmacokinetic suitability, drug-likeness and predicted safety.After over all profiles were identified,these have good binding capability with CDK2 which is protein-ligand interactions good drug-likeness characteristics and predicted oral toxicity in Class 4.Among them Carteolol was selected as the most promising candidate because of its balanced overall profile, including a docking score of −8.650 kcal/mol, 11 PLIP interactions, high predicted gastrointestinal absorption, zero Lipinski violations, zero PAINS alerts, an LD₅₀ of 810 mg/kg, and Toxicity Class 4.It can be potential repurposing candidate for targeting CDK2.Though it is suitable, need to perform including molecular dynamics simulations, CDK2 enzyme inhibition assays, binding studies and cell-based experiments to validate binding mechanism potential.

